# Characterization of Human Ectocentromeric Sites

**DOI:** 10.64898/2026.05.28.728588

**Authors:** Pasquale Saggese, Cinzia Benetti, Francesco Boccalatte, Simona Giunta

**Affiliations:** Laboratory of Genome Evolution; Department of Biology and Biotechnologies “Charles Darwin”; University of Rome “La Sapienza”; Piazzale Aldo Moro, 5 Rome 00185, Italy; Candiolo Cancer Institute, FPO - IRCCS, 10060, Candiolo, TO, Italy

**Keywords:** Ectocentromeric sites, CENP-B, CENP-B boxes, Centromere, Chromosome architecture, Gene regulation

## Abstract

Centromeres are composed of DNA repeats within chromosomes’ primary constriction. CENP-B is the only centromeric protein known to bind a specific motif, the CENP-B box, promoting kinetochore stability. We recently uncovered degenerate CENP-B binding motifs outside centromeres, whose position and orientation defines chromosome specific banding patterns. Here, we leveraged telomere-to-telomere assemblies to map conservation of these ectocentromeric sequences (ECS) across hundreds of haplotypes. We found strong negative selection acting on their occurrence along chromosome arms, implying functional constraints incompatible with stochastic drift. We classified four categories: (i) ECSs that lack CENP-B binding (∼84%); (ii) ECSs bound by CENP-B (∼10%); (iii) ECSs near CENP-B-enriched accessible chromatin (∼6%); (iv) we further identified ∼700 CENP-B binding sites outside centromeres without CENP-B boxes. Integrating chromatin conformation capture (HiC), neocentromeres and meiotic recombination mapping with CENP-B CUT&RUN, methylation and ATAC-seq data, we found heterogenous functionalities driven by distance-dependent enrichment and local contacts of boxes in inverted orientation on the same strand, analogous to ALU repeats affecting topological folding. CENP-B knockdown significantly reduced neighboring gene expression, revealing a moonlighting regulatory role outside centromeres. Our findings characterizes human *ectocentromeric sites* as evolutionarily constrained and functionally heterogeneous elements along chromosome arms with context-dependent roles in chromatin state.

**Graphical Abstract:** **Ectocentromeric sites exhibit heterogeneous CENP-B occupancy and context-dependent chromatin functions.**

Ectocentromeric sequences (ECSs) along chromosome arms fall into four categories: (i) CENP-B box motifs alone, lacking protein binding, embedded within repressed or boundary chromatin and contributing to TAD organization. Motifs with opposite orientation (forward and reverse complement) paired on the same strand may further promote self-complementary chromatin contacts analogous to Alu inverted repeats, reshaping topology with long-range looping contacts; (ii/iii) ECSs bound by CENP-B protein, associated with specific open chromatin state downstream of H3K27me3-marked compacted chromatin that modulate local accessibility and gene expression; and (iv) CENP-B binding peaks lacking a canonical box motif, located proximal to transcription start sites and linked to active gene expression. Together, ectocentromeric sites represent functionally heterogeneous elements with context-dependent roles spanning chromosome architecture, chromatin state, and transcription regulation.

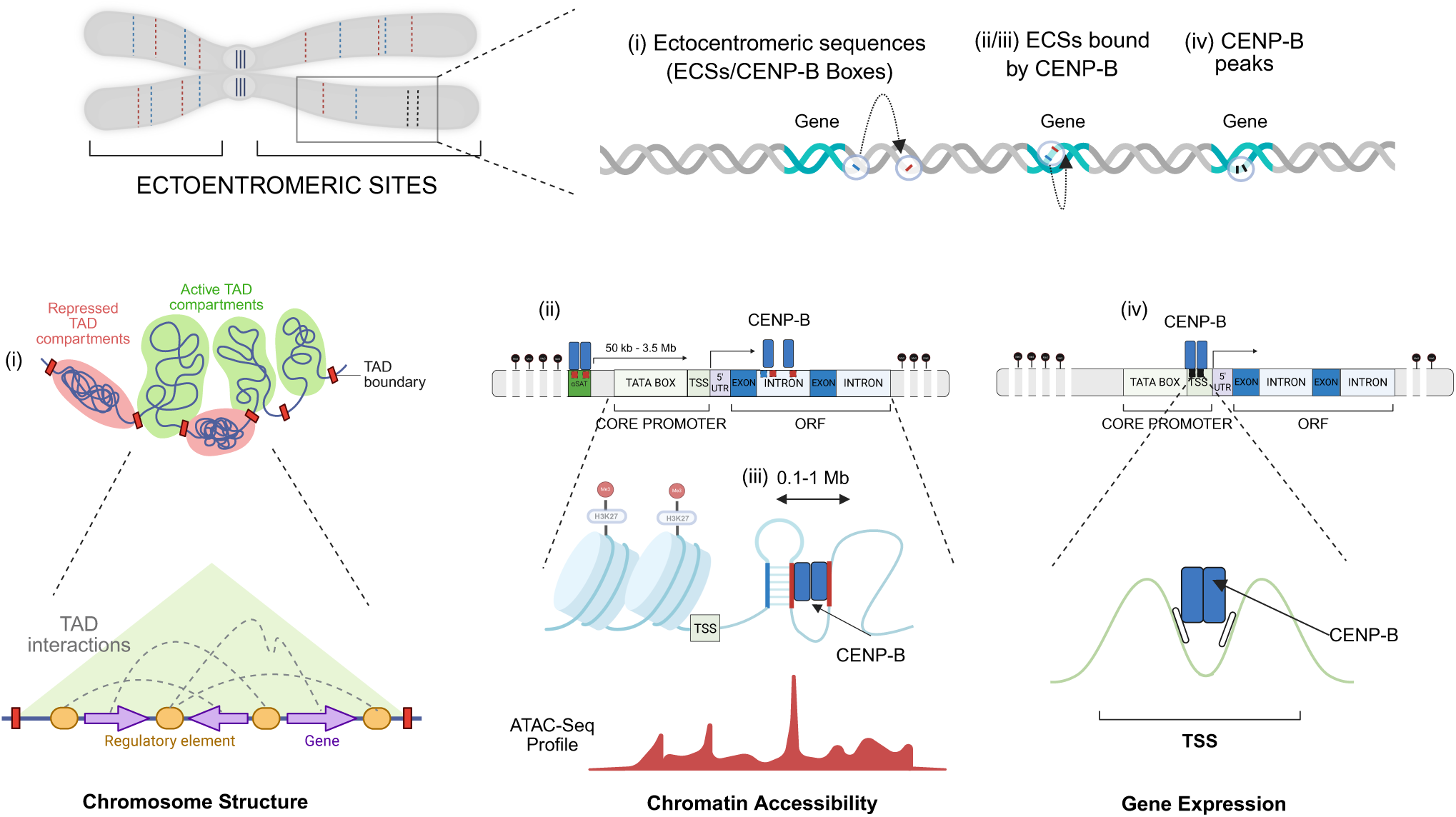

## Introduction

Repetitive DNA sequences are distributed in multiple copies throughout the genome [1] and are considered key players in genome architecture, gene regulation, and evolutionary dynamics [2,3]. Numerous studies have investigated telomeric-like sequences at internal sites, known as interstitial telomeric sequences (ITSs) [4]. ITSs were found throughout the genome away from their expected location at the chromosomes’ ends, including near centromeres and along the arms [5], and were recognized for their role in chromosomal instability [6–8]. Similar to telomeres, centromeres are regions of repetitive satellite DNA with their own genetic and epigenetic identity. In humans, they are structurally composed of alpha-satellite (αSat), where ∼170 base pairs (bp) αSat monomers are organized in tandem to make up a higher order repeat (HOR) unit, which is then reiterated over the megabase-sized centromere domain [9–12]. A small portion marks the functional region that mediates kinetochore attachment for chromosome segregation. This locus is characterized by the presence of the centromere-specific histone H3 variant centromere protein A (CENP-A) [13] in high density [14] and by dip in DNA methylation (CDR) [15]. In 1989, Masumoto et al. characterized the Centromeric Protein B (CENP-B) as the only known DNA-binding protein that specifically recognizes the CENP-B box, a 17 bp motif where only 9 bp are essential for DNA-protein contact [16]. CENP-B contributes to the centromeric organization and kinetochore assembly via direct interaction with CENP-C, thus promoting chromosome segregation [17]. Emerging research suggests that it may influence the formation of DNA loops [18] and facilitate the incorporation of histone H3.3 via DAXX chaperone, ensuring centromere chromatin integrity [19].

We recently developed a novel computational approach, the Genomic Centromere Profiling (GCP) pipeline, that enables the identification and processing of CENP-B boxes genome-wide [20]. Using this pipeline, we uncovered that each chromosome has a specific position of CENP-B boxes at a certain distance from the next, especially within the centromere, where the overall box density is ∼130-fold higher [20]. The implications of these findings are multifold, from conserved architecture in spite of underlying sequence divergence, to isolation of individual chromosome-specific centromeric reads, color-coded centromere characterization and comparison based on distance values between CENP-B boxes instead of the underlying sequence (https://github.com/GiuntaLab/GCP-Centeny). We also found occurrences of CENP-B boxes along the chromosome arms and named these “ectocentromeric sequences” (ECSs), whose position and orientation are non-random, collectively defining a chromosome-specific *centeny* map that can be used for synteny-like genomic comparison [20]. This discovery prompted us to perform a systematic characterization of ECSs across the human genome, addressing their conservation between individuals and primate species, the evolutionary constraints shaping their genomic organization, their capacity to recruit CENP-B outside the centromere, and the functional consequences of such binding on local gene expression and chromatin organization. Here, we present the results of such comprehensive analyses that integrate hundreds of near telomere-to-telomere (T2T) assemblies, CENP-B Cleavage Under Targets and Release Using Nuclease (CUT&RUN), Assay for Transposase-Accessible Chromatin with sequencing (ATAC-seq), and High-throughput Chromosome Conformation Capture (Hi-C) datasets to answer these questions, providing the first definition and functional classification of ectocentromeric sites.

## Methods

### CENP-B Box annotation

To query a selected reference genome for the CENP-B Box sequence, the *fuzznuc* (EMBOSS v6.6.0.0) program [21] has been implemented in the GCP tool.

The query used for all the analyses includes the CENP-B box DNA motif, which contains the specific 9 nucleotides essential for CENP-B protein binding, embedded within a larger query of 17 nucleotides (*TTCG****A**CGGG*). The tool searches for this motif within nucleotide sequences and reports occurrences on both the forward (+) and reverse complement (-) strands. Input sequences can be provided as FASTA files from long-read sequencing platforms (e.g., ONT or PacBio) or as FASTA files of a genome assembly at the chromosome or contig level. The output is a BED file with the following columns: chromosome or sequence ID, start position (0-based, inclusive), end position (1-based, exclusive), and strand (+ or -).

### Centeny Map

The consensus centeny maps were generated using the karyoploteR package (R/Bioconductor)[22] with CHM13 T2T v2.0 as the reference genome. Chromosomal lengths were defined directly from CHM13 T2T v2.0 coordinates as a custom GRanges object, replacing the karyoploteR default hg38 genome. For each ECS, a bar was drawn with a fixed width of 500 kb (±250 kb centered on the site position) and height proportional to the frequency of that position across the 215 analyzed HPRC/HGSVC haplotypes (0-100%). Fig. 1A includes all 1,359 ECS identified on CHM13 T2T v2.0, with bar color indicating motif orientation: blue (#2563EB) for forward (+) (n=601) and red (#DC2626) for reverse complement (-) (n=758); the 505 positions with frequency =0% are shown as zero-height bars. Fig. S1 includes only the 854 ECS with frequency >0%, classified into two categories displayed with distinct colors: High-frequency ECSs (frequency >=75%, n=18, blue #1565c0) and Polymorphic ECSs (0% < frequency <75%, n=836, orange #ff9800); Bars for polymorphic ECSs were plotted before those for conserved ECSs to ensure the visibility of highly conserved sites.

**Figure 1.**
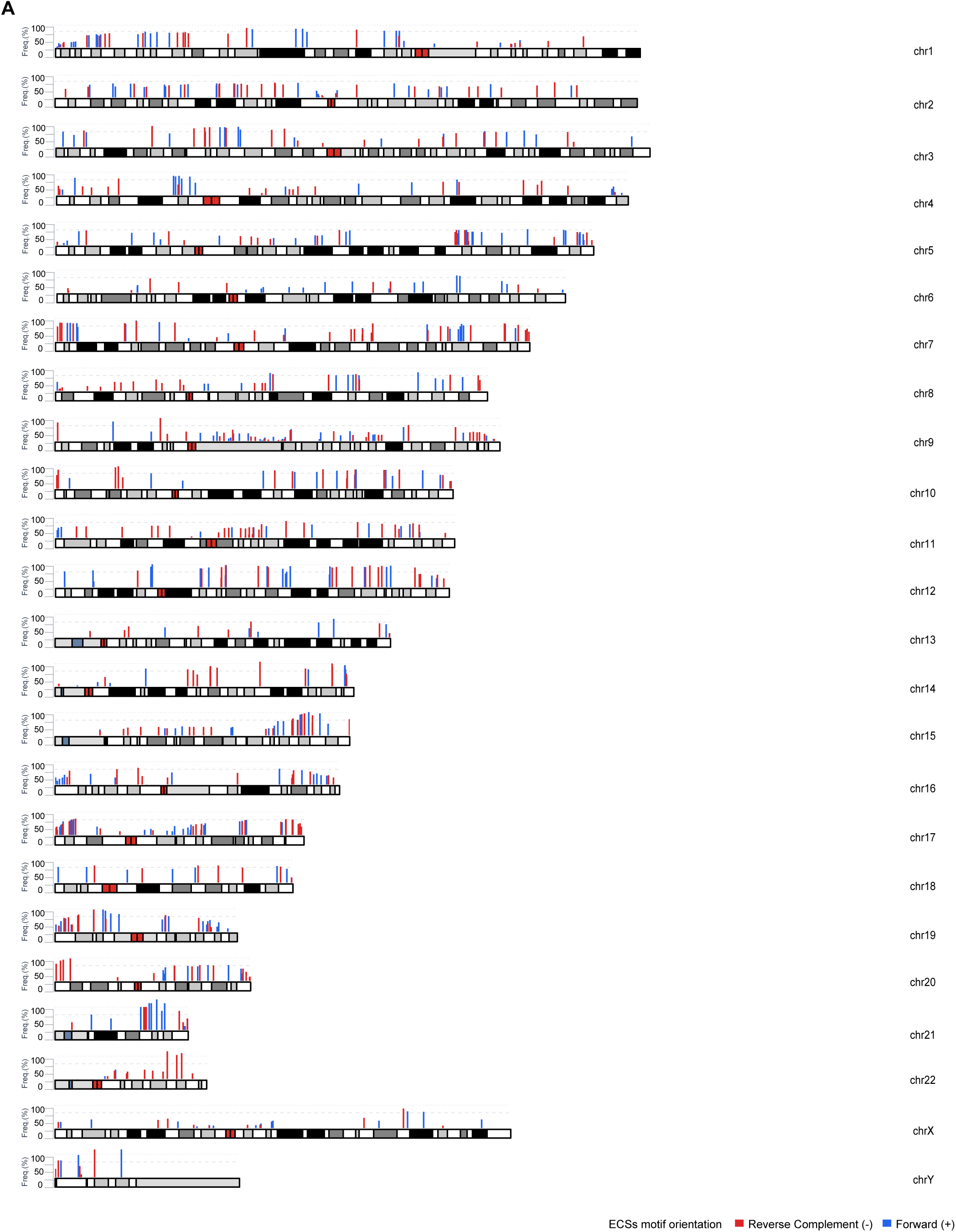
Consensus Human Centeny Map derived from multiple individuals’ genomes. A) Karyotype representation of all 1,359 ectocentromeric sequences (ECSs) identified on CHM13v2.0. Bar height indicates ECSs frequency across 215 HPRC/HGSVC assemblies (0-100%). Blue = forward (+) (n=601); red = reverse complement (-) (n=758). Of the 1,359 ECS, 854 (62.8%) are present in at least one genome; 505 (37.2%) are absent in all analyzed genomes (bar height = 0), of which 322 overlap alpha-satellite regions and 183 lie outside alpha-satellite arrays.

### ECS conservation frequency analysis

To assess ECSs conservation across the human population, the 1,359 ECS positions identified on CHM13v2.0 were used as a reference and their presence evaluated across 215 phased haplotypes from 109 individuals of the HPRC and HGSVC consortia. For each assembly, ECSs were remapped to CHM13 coordinates using a nearest-neighbor, strand-specific approach. For each ECS, the closest CHM13 position on the same strand with start_CHM13 < start_ECS was identified within a chromosome-specific distance threshold. In cases of multiple assignments, collisions were resolved by retaining the closest ECS. The frequency of each CHM13 ECSs position was then calculated as the proportion of the 215 haplotypes in which that position was covered by a remapped ECS (range: 0–100%). Visual analysis was performed to assess ECS conservation across T2T reference genomes. Both datasets were then compared and plotted in a single bar plot. Genome-wide, ECSs positions were classified into three categories based on their frequency: High-Frequency ECSs (frequency ≥75%, n=18), Polymorphic ECSs (0% < frequency <75%, n=836), and CHM13-specific ECSs (frequency =0%, n=505). The proportion of each category is shown in a horizontal bar chart. The distribution of High-Frequency and Polymorphic ECSs positions per chromosome was visualized as a stacked bar chart showing the proportion of each frequency subcategory (Conserved 100%, High 75–99%, Medium 50–74%, Variable 25–49%, Rare <25%) for each of the 24 chromosomes. The 505 CHM13-specific ECSs positions were excluded from the stacked bar chart and analyzed separately by intersection with CHM13 alpha-satellite annotations from HumAS-HMMER_for_AnVIL using coordinate-based overlap, classifying them as inside αSat (n=322, 63.8%) or outside αSat (n=183, 36.2%). Three additional supplementary analyses were performed to characterize non-random patterns in ECSs distribution. First, the correlation between distance from the centromeric border and ECS frequency was calculated using the Spearman coefficient, with ECS grouped into quartiles of centromeric distance for visualization. Second, ECSs were assigned to chromosomal arms (p or q) using centromeric border coordinates and median frequency was compared between arms using the Mann-Whitney U test. Third, motif orientation bias was assessed for each chromosome and genome-wide using a chi-square test against the expected 50/50 ratio. All p-values are two-sided.

### CENP-B box motif analysis and integrity scoring

To characterize the sequence properties of the CENP-B box motif in highly conserved ECSs, two complementary approaches were used. First, genomic sequences of 17 bp were extracted for all 1,359 ECS positions from the CHM13v2.0 reference genome using the Biostrings package (R/Bioconductor) [23]. For sequences on the negative strand (n = 758), the reverse complement was computed to orient all sequences in the 5′→3′ direction of the CENP-B box motif. The 18 ECSs with frequency ≥75% were subjected to de novo motif discovery using MEME Suite (v5.5.9; https://meme-suite.org), with motif width fixed at 17 bp (min = max = 17), a one occurrence per sequence model (oops), and the search limited to a single motif. The resulting consensus motif was compared position-by-position to the canonical CENP-B box sequence YTTCGTTGGAARCGGGA [16,24] (Y = C/T, R = A/G). Sequence logos were generated using the ggseqlogo package (R). Motif integrity was quantified using a Motif Integrity Score (MIS), defined as the number of matches to the 9 fixed nucleotides of the canonical CENP-B box (positions 2, 3, 4, 5, 10, 13, 14, 15, 16; expected bases: T, T, C, G, A, C, G, G, G). For each ECS, MIS was calculated as the number of matching positions (range: 0–11), using IUPAC-aware base comparison. MIS was calculated separately for four groups: High-Frequency ECSs (frequency ≥75%, n=18), Polymorphic ECSs (0% < frequency <75%, n=836), CHM13-specific ECSs inside αSat regions (frequency =0% and overlapping alpha-satellite, n=322), and outside αSat regions (frequency =0% and outside alpha-satellite, n=183). Group differences were assessed with the Kruskal-Wallis test followed by pairwise Wilcoxon tests with Bonferroni correction. All analyses were performed in R (Version 4.5.0).

### Secondary structure propensity analysis (Minimum Free Energy, MFE)

Consecutive ECSs pairs in the 100 kb – 1 Mb range were identified in both haplotypes of the RPE-1v1.1 genome (n = 448 motifs in haplotype 1, n = 440 in haplotype 2). For each motif, a ±250 bp window centered on the motif was extracted using bedtools getfasta. Sequences were converted from DNA to RNA and submitted to Minimum Free Energy (MFE) computation using RNAfold from the ViennaRNA Package (v2.7.2) [25] with default parameters at 37 °C. More negative MFE values indicate a greater intrinsic propensity for stable secondary structure formation. Two types of control sequences were generated. Three scrambled controls were produced by progressively permuting the CENP-B motif, as described in section 1.2, and identifying genomic matches of each permuted motif (Scrambled 1: n = 342 / 362; Scrambled 2: n = 888 / 840; Scrambled 3: n = 1110 / 1062, for haplotype 1 / haplotype 2, respectively). An additional dinucleotide-preserving control (Shuffled) was generated by permuting each CENP-B box sequence while preserving its dinucleotide frequency, with 10 independent replicates per sequence (set.seed = 42; n = 4480 / 4400), thereby controlling nucleotide-composition effects on structure propensity. The number of motifs differs across scrambled controls owing to the intrinsic genomic rarity of permuted motifs. The Wilcoxon rank-sum test is robust to unequal sample sizes and does not require down sampling. Statistical comparisons between CENP-B box motifs and each control were performed using the two-sided Wilcoxon rank-sum test. As an orientation control, MFE values were additionally stratified by the relative orientation of the two motifs within each pair (Same vs Opposite) and compared using the two-sided Wilcoxon rank-sum test, linking MFE values to the corresponding sequences based on genomic position and motif identifiers.

### ECSs data processing

ECS counts and motif analyses were performed using the BED files generated by the GCP pipeline for both the CHM13v2 and RPE-1v1.1 reference genomes. The simulation analysis was performed by querying the selected scrambled and the αConsensus sequences using the *fuzznuc* (EMBOSS v6.6.0.0) program. Distance values between consecutive CENP-B boxes were calculated as the difference between the start coordinate of a given occurrence and the end coordinate of the preceding occurrence.

Distance values obtained from the scrambled sequences were randomly down sampled to match the number of values in the original dataset, allowing for direct comparison. Data visualization was performed using GraphPad Prism (v10.0.0; GraphPad Software) and *ggplot2* (v3.5.1) [26] in R (v4.5.0) to generate plots for genome-wide and chromosome-level analyses. The analysis was performed both genome-wide and on ECSs located at least 5 Mb away from centromeres.

### Annotations and Functional Relevance of ECSs

To characterize ectocentromeric sites (ECSs) across the RPE-1v1.1 reference genome, we used the BED files previously generated from the analysis of selected publicly available datasets: CUT&RUN for CENP-B (GSE132193) [27], ATAC-seq (GSE209659) [28], and H3K27me3 ChIP-seq (GSE184030) [29], all performed in hTERT RPE-1 cells. For the CENP-B CUT&RUN dataset, we downloaded three IP replicates (SRR9201837, SRR9201839, SRR9201844) and three matched input controls (SRR9201838, SRR9201846, SRR9201847). CENP-B peaks were defined and analyzed as broad enrichment regions. For visualization purposes, bigWig tracks were generated by normalizing the CENP-B signal to the matched input control and were inspected using the Integrative Genomics Viewer (IGV).

### RPE-1v1.1 gene annotation

Liftoff (v1.6.3) was used to map genes from the ENSEMBL GRCh38.p14 annotation file (version 112) to the RPE-1 genome. The annotation files for Hap1 and Hap2 were used to intersect with CENP-B box occurrences, ATAC-seq, and CENP-B protein peaks.

### Analysis of ECS orientation bias

To assess the orientation bias of ECSs, we counted the number of motifs in forward and reverse-complement orientation for each genome assembly and compared them to a null hypothesis of equal probability (p = 0.5) using a two-sided binomial test implemented in R (Version 4.5.0).

### Kolmogorov–Smirnov (K-S) test for ECS spacing

To assess whether the genome-wide distribution of distances between consecutive ECSs differs from that expected by random sequence organization, we performed two-sample Kolmogorov–Smirnov (K-S) tests. For each genome assembly, ECS distance distributions were compared to those derived from scrambled or αSat-derived control sequences of equal length. Tests were performed using the “ks.test” function in R (version 4.5.0), and significance was assessed based on the D statistic and corresponding p-values. In cases where tied values occurred, p-values were reported as approximate.

### Hi-C Analysis

Each RPE-1v1.1 haplotype was treated as an independent reference genome, indexed separately, and restriction fragments corresponding to the Arima Hi-C protocol were predicted using the *HiC-Pro* generate_site_positions.py utility [30]. Hi-C reads were then aligned, filtered, and deduplicated independently against each haplotype using the *HiC-Pro* pipeline with default parameters [30]. Lane-merged, deduplicated valid read pairs were used as input for all downstream analyses within the *HiC-Bench* framework [31]. Haplotype-resolved Hi-C contact matrices were analyzed using the *HiC-Bench* matrix-sparse and virtual4C pipelines [31]. ECS loci were used as viewpoints, and interaction profiles were computed in a rolling-window fashion. Specifically, for each viewpoint, interactions were aggregated in 200-bp bins by summing valid read pairs within a ±5 kb local window, extending up to ±2.5 Mb along the chromosome. Interaction profiles were normalized to counts per million (CPM) by dividing by the total number of valid read pairs per sample and exported in bedGraph format for downstream visualization and comparison. Topologically associated domains were identified using the *HiC-Bench* domain module on ICE-normalized and filtered contact matrices at 20 kb resolution [31,32]. TAD boundaries were extracted from both haplotypes and combined for downstream analyses. Boundary coordinates were defined as single-base regions corresponding to TAD start and end positions. Contact matrices were accessed directly from .hic files in R using the strawr package [33] with raw (NONE) normalization. Chromosome name conversion between BED format (chrN_hapX) and Hi-C format (CM116*) was performed using a mapping table. Distances from each ECS locus to the nearest TAD boundary were computed using GenomicRanges [34]. ECS loci overlapping annotated centromeric or pericentromeric regions (±5 Mb) were excluded. Statistical significance was assessed using a permutation test (1,000 iterations), in which ECS positions were randomized along their respective chromosomes while preserving chromosomal distribution and excluding centromeric regions. Observed median distances were compared to the distribution of permuted medians to derive empirical p-values.

Local insulation scores were computed at 25 kb resolution for all ECS loci and for matched scrambled control positions. For each locus, a symmetric contact window spanning ±10 bins (250 kb on each side) was extracted. The insulation score was defined as log₂(cross / √(left × right)), where cross represents contacts spanning the two sides of the central bin, and left and right represent within-domain contacts on each side. More negative values indicate reduced local chromatin interconnectivity, consistent with chromatin domain boundaries. ECS-associated values were compared to those of three independent scrambled control sets sharing the same chromosomal distribution using two-sided Wilcoxon rank-sum tests. A/B compartments were defined from the first principal component (PC1) of normalized Hi-C contact matrices, with genomic bins assigned to compartment A or B based on PC1 sign. ECS loci were classified into mutually exclusive functional categories (box-only, protein-only, box-protein and box-near-protein) based on the presence of the canonical CENP-B box motif and CENP-B binding detected by CUT&RUN. ECS coordinates were intersected with compartment annotations using GenomicRanges::findOverlaps (select = “first”). Deviations from the expected 50/50 A/B distribution were assessed using binomial tests. Visualization. Representative Hi-C contact maps were visualized using Juicebox [35].

### Aggregate Peak Analysis (APA) of ECS pairs

To test whether pairs of nearby ECS loci engage in preferential Hi-C contacts, we performed distance-stratified Aggregate Peak Analysis (APA) on two ECS categories: Box-only (BO) and Box-near-protein (BNP). Haplotype-resolved Hi-C contact maps (.hic files) were accessed in R using the strawr package [33] at 25 kb resolution with raw (NONE) normalization.

All ECSs coordinates were pre-filtered with a using 5 Mb cutoff from pericentromeric to exclude alpha-satellite regions, and chromosome names were converted from BED format (chrN_hapX) to Hi-C format (CM116*) via a mapping table. For Box-near-protein (BNP), each row of the annotation file represents a biologically defined pair consisting of CENP-B protein site and its adjacent CENP-B box (n = 93 pairs in haplotype 1, n = 93 in haplotype 2; distance range 3.2–995 kb). CENP-B position was used as the anchor for matrix extraction.

For Box-only (BO), single CENP-B box sites (n = 748 in haplotype 1, n = 725 in haplotype 2) were sorted by genomic position within each chromosome and consecutive nearest-neighbor pairs were constructed (box i with box i+1). Box-only pairs were restricted to the 0–1 Mb distance range for analysis (n = 278 in haplotype 1, n = 277 in haplotype 2; 555 pairs total).

### Distance-matched random controls

For each real pair, five random control pairs were generated on the same chromosome, preserving the exact inter-element distance and orientation of the real pair, with random positions sampled uniformly within the chromosome length (set.seed = 42 for reproducibility). APA matrices (11 × 11 bins) were extracted for each real and random pair, centered on the predicted contact point and spanning ±5 bins (±125 kb at 25 kb resolution).

Pairs falling in regions of insufficient Hi-C coverage were discarded (retention 91–95%), yielding 254/265 (haplotype 1/2) real BO matrices, 87/88 real BNP matrices, and the corresponding distance-matched random matrices (∼5× the real counts). Pairs were stratified into 10 distance bins of 100 kb each (from 0–100 to 900–1000 kb). For each bin, the central matrix value [6,6], corresponding to the direct Hi-C contact between the two paired elements, was extracted from real and random matrices. Real versus random central values were compared using the Wilcoxon rank-sum test (two-sided for Box-only, where both depletion and enrichment were a priori possible; one-sided ‘greater’ for Box-near-protein, testing the enrichment hypothesis).

Effect magnitude was quantified as the fold-change (ratio of median real to median random central values) and as Cohen’s d (standardized mean difference using the pooled standard deviation; |d| < 0.2 negligible, 0.2–0.5 small, 0.5–0.8 medium, > 0.8 large). To confirm that the effects were independent of haplotype-specific genome assembly, the focal distance bins (Box-only: 0–100 kb; Box-near-protein: 100–200 kb) were re-analyzed separately for haplotype 1 and haplotype 2, using only the real and random pairs of each haplotype against the corresponding Hi-C contact map (no pooling). Concordance was assessed by three criteria: (i) Wilcoxon p < 0.05 in both haplotypes; (ii) consistent sign of Cohen’s d; and (iii) consistent direction of the fold-change.

Aggregate APA matrices were rendered as divergent heatmaps (blue–white–red), with the color scale calibrated independently for each distance bin to account for the intrinsic distance-dependent decay of Hi-C contacts. All analyses were performed in R using strawr [33], dplyr [36], and ggplot2/patchwork [26].

### Spatial overlap and permutation analysis between ECSs and DMC1 peaks

Genomic coordinates of ectocentromeric sequences (ECSs) were intersected with publicly available DMC1 binding sites obtained from SSD-Seq experiments [37]. Overlaps were defined as any base-pair intersection between an ECS interval and a DMC1 peak.

To evaluate whether the observed overlaps exceeded random expectation, we performed permutation tests that preserve both chromosome identity and interval length. ECS coordinates were randomly repositioned within the genomic space under investigation using a custom randomization procedure, and the number of intersections with DMC1 peaks was recalculated at each iteration. Two independent permutation frameworks were applied: PAR-restricted analysis, in which ECSs located within pseudoautosomal regions were shuffled exclusively within PAR coordinates; non-PAR analysis, in which ECSs outside PARs were randomized only across the remaining genomic regions. For each condition, 10,000 permutations were performed to generate a null distribution of expected overlaps. Empirical p-values were calculated as the fraction of permutations yielding overlap counts equal to or greater than the observed value. Fold enrichment was computed as the ratio between observed and mean expected overlaps, and z-scores were derived from the permutation distributions.

### qPCR Analysis

hTERT RPE-1 cell line was cultured in DMEM/F12 medium supplemented with 10% fetal bovine serum (Gibco, 10270-106), 100 μg/ml streptomycin, 100 U/ml penicillin (Euroclone, ECB3001D), and 2 mM glutamine (Euroclone ECB3000D). Cells were grown at 37 °C in a humidified atmosphere of 5% CO2 and tested negative for mycoplasma contamination.

The following siRNAs targeting CENP-B (siControl: AATTCTCCGAACGTGTCACGT; siRNA-1: CGCCTTGAGAAGCACAGTTTA; siRNA-2: CCGGGAGAAGTCACGGATCAT; siRNA-3: CACGCTGAGCACGATCCTGAA; siRNA-4: TTCGAGGTCTTGGACATCAAA) were purchased from Qiagen (Cat. No. 1027415). All four siRNAs were tested for efficiency, and the two yielding the strongest CENP-B knockdown were selected. Transfection was performed according to the manufacturer’s instructions using Lipofectamine RNAiMAX (Thermo Fisher, Cat. No. 13778-075). The following primers were used for the qPCR analysis: CENP-B (Forward: GCGCTTGTTCTTCAGGATCG, Reverse: CGGGAGAAGTCACGGATCAT), ALG10B (Forward: CTGTTTGACTTGGCCCTACA, Reverse: GACTACTCCGATCGCCAATAAC), PRDM16 (Forward: CCACATCGCTGCTCAAGA, Reverse: TGGCTGCTGTTGCTGAC), CENP-C (Forward: TTCCAGTGAGTCCAAGAACAAA, Reverse: TTACCACCCACCAATCAGATG), and GAPDH (Forward: GTCAACGGATTTGGTCGTATTG, Reverse: TGTAGTTGAGGTCAATGAAGGG). All primers were designed by NCBI Primer-BLAST (RRID: SCR_003095) and purchased from Integrated DNA Technologies. mRNA levels were normalized to GAPDH (ΔCt Ct gene of interest - Ct GAPDH) and presented as relative mRNA expression (2ΔCt). The experiment was performed in biological triplicate, and data were pooled for the analysis.

## Results

### 1.1 Conservation and Evolution of ECSs

Leveraging the T2T human reference genome CHM13v2 [38] and the near-complete assembly of the hTERT RPE-1 cell line recently generated by our laboratory [14], we previously discovered that the overall chromosomal organization of ECSs, including their positional patterns and orientation, is preserved between individuals [20]. The conserved position and specific orientation of CENP-B boxes across human chromosomes define the human *centeny* map – a computational banding framework for chromosome recognition and characterization [20].

To assess the exact conservation of each ectocentromeric sequence and to draft a consensus *centeny* map as representative as possible of the general population, we leveraged cohorts of currently available assemblies from Human Pangenome Reference Consortium (HPRC) [39], and Human Genome Structural Variant Consortium (HGSVC) [40] (Fig. 1A). Using the ECSs mapped on CHM13v2.0 as a reference, we assessed their conservation across 215 assemblies of phased haplotypes from both consortia. Beyond positional and orientation data, the resulting consensus *centeny* map integrates ECS frequency across genomes (Fig. 1A), revealing a substantial variability driven by inter-individual differences in occurrences. We classified only 18 sites (1.3%) as high-frequency ECSs (frequency ≥75%), while 836 positions (61.5%) correspond to polymorphic ECSs (1% < freq < 75%) and 505 positions (37.2%) represent CHM13-specific ECSs (frequency = 0%) absent in all other analyzed individuals (Supp. Tables 1-2).

When stratified by chromosome, ECS conservation patterns varied markedly (Fig. 2B). ECSs with variable frequency (25-49%) were observed across most chromosomes, reflecting the overall low conservation of ECSs in the human population, with chromosomes 5 and 11 showing the highest proportion. Rare ECSs (<25%) were also widely distributed, with notable enrichment on chromosome 9, 22 and X, consistent with the elevated structural variability in their pericentromeric regions. ECSs with intermediate conservation (50-74%) were present on most chromosomes, with a relative enrichment on chromosomes 10 and 12. In contrast, highly conserved ECSs (≥75%) were restricted to a limited number of loci, primarily on chromosomes 21 and Y, while fully conserved ECSs (100%) were detected exclusively on chromosome 21. A genome-wide conservation map (Fig. S1) further indicates that ECS conservation is partially heterogenous across individuals, with a fraction of loci showing high population frequency, while the majority display variable frequencies. Highly frequent ECSs are restricted to a limited number of chromosomes, forming distinct hotspots, whereas intermediate-frequency ECSs are broadly distributed but heterogeneous. 505 ECSs were classified as absent (frequency = 0) due to a distinct genomic distribution that were visually challenging to overlap. Of these, 322 (63.8%) localize within pericentromeric or peripheral alpha-satellite regions of CHM13, consistent with extensive structural variability of high order repeat (HOR) arrays between individuals. The remaining 183 positions (36.2%) were located outside the alpha-satellite regions. Interestingly, we found a significant positional gradient with respect to centromere distance (Spearman ρ = 0.209, p = 1.2 × 10⁻¹⁰). ECSs located closer to the centromere (<8.2 Mb) showed lower conservation (median frequency 26%) compared to those in distal regions (>45.5 Mb; median 42.5%) (Fig. S4B). This pattern indicates that pericentromeric ECSs are more variable across individuals, consistent with the elevated structural instability and dynamic evolution of these genomic regions, or may reflect difficulties in correctly assembling these regions. Indeed, comparison between chromosome arms showed that ECSs located on the p arm exhibit significantly higher conservation than those on the q arm (median frequency 39.0% vs 31.9%, Mann–Whitney U test, p = 0.005). However, acrocentric chromosomes are largely depleted of ECSs on the p arm (Fig. S4C). In addition, ECSs displayed a significant genome-wide orientation bias toward the negative strand (55.8% vs 44.2%, chi-square test, p<0.0001), with chromosome-specific patterns. In particular, chromosomes 6 and 14 showed an enrichment of ECSs on the positive strand, while chromosomes 9, 11, and 20 were predominantly on the negative strand (Fig. S4D), likely driven by chromosome-specific differences in pericentromeric structure and evolutionary dynamics.

**Figure 2.**
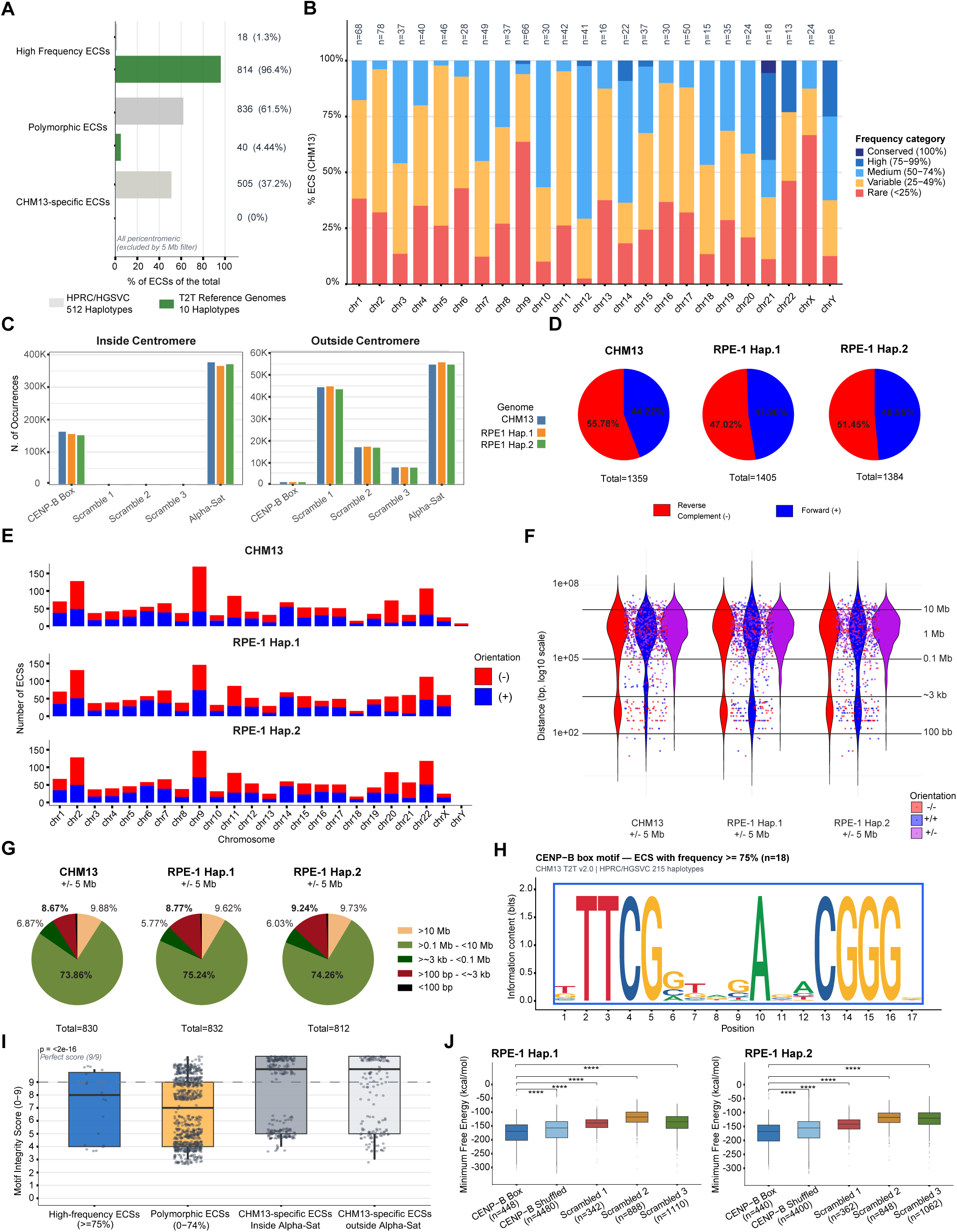
Genomic Distribution and Structural Features of the ECSs. A) Bar plot of the distribution of 1,359 ECSs classified by conservation: High-frequency (freq ≥ 75%, blue), polymorphic ECSs (1% < freq < 75%, orange), and CHM13-specific ECSs (freq = 0%, grey). B)Stacked bar chart showing the distribution of 854 ECSs with frequency >0% by conservation category: Conserved (100%, dark blue), High (75-99%, blue), Medium (50-74%, light blue), Variable (25-49%, orange), Rare (<25%, red). n = ECS per chromosome with at least one remapped sample. C) Bar plot illustrates the total counts of CENP-B Box, scrambled sequences, and αConsensus, either inside or outside the centromere, in both CHM13 and RPE- 1 reference genomes. D) Pie charts representing the genome-wide total counts and the percentage of forward and reverse complement of ECSs in CHM13 and RPE-1 haplotypes 1 and 2. For the analysis, we considered the occurrences within the divergent αSat arrays located outside the centromere, provided by HumAS-HMMER_for_AnVIL. E) Stacked bar plots showing the total number of forward and reverse complement ECSs in each chromosome. F) Violin plot of whole-genome ECS distances (log2 scale) in CHM13 and RPE-1 haplotypes, following application of the 5Mb cut-off. G) Pie charts showing the percentage of the CENP-B box distances (bp) in selected ranges in CHM13 and RPE-1 haplotypes. H) sequence logo of the CENP-B box motif identified by MEME analysis in the 18 ECS positions with frequency >= 75% across 215 HPRC/HGSVC haplotypes. The motif consensus is KTTCGGKDGAGACGGGR (E-value = 3.7x10-73). Fixed positions (2-4, 5, 13-16) show maximum information content. Degenerate N positions show non-random nucleotide preferences, particularly G at positions 6 and 9, A at positions 10 and 12, and purine bias at positions 1 and 17. Sequences from negative-strand ECS (n=10) were reverse-complemented prior to analysis. I) Box plots showing the distribution of Motif Integrity Score (MIS) across four ECS groups: High-Frequency ECSs (frequency ≥75%, n=18), Polymorphic ECSs (0% < frequency <75%, n=836), CHM13-specific ECSs inside αSat arrays (frequency =0% and overlapping alpha-satellite, n=322), and outside αSat arrays (frequency =0% and outside alpha-satellite, n=183). MIS is defined as the number of matches at the 9 fixed core positions of the canonical CENP-B box sequence YTTCGTTGGAARCGGGA (range: 0–11). Individual data points are shown as jittered dots. Statistical significance was assessed by Kruskal-Wallis test followed by pairwise Wilcoxon tests with Bonferroni correction; significance levels are indicated above each comparison. The dashed line indicates a perfect core match score (9/9). J) Boxplot of Minimum Free Energy (MFE) distributions for sequences flanking ECS motifs compared to controls.

To better understand whether heterogeneity in occurrence of ECSs was influenced by accuracy during genome reconstruction, we focused on currently available diploid human references, T2T human genomes and chromosome-level assemblies, including HG002 maternal and paternal [41,42], YAO maternal and paternal [43], H9 maternal and paternal [44], I002C maternal and paternal [45], and RPE-1v1.1 haplotypes 1 and 2 [14] (Fig. S2). Qualitative analysis of diploid T2T human genomes revealed largely consistent ECS patterns along both p and q arms across all assemblies examined. Although a small number of ECSs occupy non-canonical positions, these cases are rare and do not disrupt the overall chromosomal organization. Visual analysis of ECS positions across T2T reference genomes (Fig. 2A) revealed that the proportion of ECSs classified as high-frequency is markedly higher than that observed within haplotypes from consortia (96.4% vs. 1.3%), whereas polymorphic ECSs are less represented in T2T genomes. These data indicate that, despite the variability observed for individual ECSs, their global organization remains highly conserved. At the same time, ECSs exhibit individual-specific patterns, suggesting a precise and potentially functional role. To strengthen the evidence for ECS conservation and assess their evolution, we expanded our analysis by benchmarking the human *centeny* map generated from the CHM13v2 against the latest T2T genome assemblies obtained from different primate species, including *Pan troglodytes* (chimpanzee), *Pan paniscus* (pygmy chimpanzee), *Gorilla gorilla gorilla* (western lowland gorilla), and *Sumatran orangutan* (pongo abelii) [46] (Fig. S3) Since 1:1 chromosome comparisons are complicated by karyotypic rearrangements such as fusions and fissions that may exist in different species. Therefore, our final goal was to assess whether clusters of ECSs exhibit a conserved organization indicative of evolutionary constraint. Indeed, the “inter-species *centeny* map” generated for a visual comparison revealed that the overall ECSs pattern is largely conserved across primate species (Fig. S3), with particularly high similarity between the two *Pan* assemblies as expected due to evolutionarily proximity to humans. In particular, the conservation of ECS organization on the X chromosome between humans and chimpanzees is consistent with its well-known structural and sequence similarity across primates [46,47], suggesting that ECS organization reflects overall sequence conservation and evolutionary constraints. In contrast, a limited similarity was detected in the gorilla and orangutan genomes. However, small ECS clusters resembling the human pattern were identified, especially in the chimpanzee and bonobo genomes on chromosomes 3, 4, 9, 10, and 15. These results imply that the consensus *centeny* map, similarly to chromosome banding dictated by heterochromatin and chromosome anatomy, is largely human-specific, highlighting its limitation for cross-species applications. Collectively, these findings suggest that, ECS positions and overall chromosomal organization is largely conserved within humans when assessed in chromosome-level assemblies, and only partially conserved across primates, supporting the use of the consensus *centeny* map as a species-level reference for chromosome comparison and characterization. Because conservation is near-absolute across 8 complete assemblies, while variation was observed when expanding to over 200 haplotypes, these analyes likely captures a mixture of true evolutionary diversity, haplotype-specific polymorphism – especially in specific repeat-rich regions such as pericentromere, as well as assembly errors and alignment uncertainty using *centeny* maps on contigs-level genomes.

### 1.2 Selection of CENP-B boxes on chromosome arms

The CENP-B box is a 17 bp motif. However, *in vitro* binding studies have shown that only 9 base pairs are required to make contact with the CENP-B protein for its recruitment [16]. Hence, the string query for CENP-B boxes contains 8 degenerate positions out of 17. Once we established that ECSs occupy conserved genomic positions across individuals, we next asked whether their presence along chromosome arms reflects functional retention under selective pressure or merely the random occurrence. To address this, we used the GCP pipeline [20] to compare the occurrence of CENP-B boxes (5’ NTTCGNNNNANNCGGGN 3’) with that of three other 17-mers sharing specific features of the CENPB box, as follows: (1) Scramble-1 (5’ TNCTNGNNANNCNGGNG 3’), generated by pairwise switching of nucleotides along the motif, thereby preserving both the number and type of fixed nucleotides of the CENP-B box motif used to identify the ECSs; (2) Scramble-2 (5’ NCAGTNNNNGNNTACAN 3’) was created by permuting only the defined (non-degenerate) nucleotides while maintaining the position of the degenerate bases; (3) Scramble-3 (5’ NGGGCNNNNANNGTTCN 3’) was obtained by switching and flipping the last 5 nucleotides at both ends of the motif, while leaving the core sequence unchanged; (4) additionally, we used another αSat consensus sequence “αConsensus” present in the centromeric monomers just upstream of the CENP-B box. All queries contained the same number of degenerate bases, yet their occurrence varied widely. As expected, the scrambled motifs were underrepresented in the centromeric sequences compared to the CENP-B box and αConsensus (Fig. 2C), indicating the strong selection for specific DNA within the centromere. Yet, CENP-B box occurrence is lower than all other motifs and below expectations based on random occurrence of such a short sequence along the genome (Fig. 2C). Strikingly, CENP-B boxes were ∼30-fold less frequent than scrambled controls and ∼40-fold less than the αSat consensus sequence along chromosome arms (Fig. S4E), demonstrating active evolutionary constraint reducing the occurrences of ECSs outside centromere, and thus pointing away from random occurrences due to sequence drift and more likely to a functional preservation under selective pressure.

### 1.3 Orientation and distribution of ECSs along chromosome arms

The presence and conservation of ECSs across genomes, including primate species, point to a potential role in maintaining structural integrity or chromatin organization. For this reason, we set out to characterize their genomic nature. First, we analyzed the number and the orientation of ECS motifs along all chromosomes, considering both the forward and reverse complement sequences found within the same strand (Supp. Table 3-5). We identified a total of 1359 ECSs in the CHM13v2 genome, and 1405 and 1384 in RPE-1v1.1 haplotypes 1 and 2 respectively, with a slight bias toward the reverse complement orientation in both reference genomes (Fig. 2D). Notably, this orientation does not reflect “strandness” but rather a non-random alternation of forward and reverse complement elements along chromosome arms. To assess whether ECS orientation deviates from random distribution, we applied a binomial test based on Chargaff’s second parity rule [48,49], which predicts equal frequencies of forward and reverse complement motifs on the same strand. Our analysis revealed that the CHM13v2 shows significant deviation from random orientation at the genome-wide level, while RPE-1v1.1 haplotypes follow parity expectations (Supp. Table 6), suggesting that orientation biases in ECS may be influenced by local genomic features rather than intrinsic properties of the ECS motifs themselves.

We next investigated whether these orientation biases are uniformly distributed across the genome or driven by specific chromosomal regions. The analysis at the chromosome level showed that in both genomes, ∼70% of chromosomes did not exhibit significant deviation from random orientation, while the remaining ∼30% showed local orientation biases (Fig. 2E, Supp. Table 6), indicating that the observed genome-wide orientation bias is primarily driven by a limited number of chromosomes exhibiting local heterogeneity, rather than being a widespread feature of ECS motifs. We wondered whether the presence of alphoid DNA repeats clusters in the chromosome arms, described as containing non-functional CENP-B boxes compared to the canonical centromeric αSat arrays [15], affected our analyses because it took into consideration the ECSs located in proximity to the centromere, closely spaced presumably due to the presence of repetitive DNA. To assess whether these regions could influence the distribution of the ECS motif orientation along chromosome arms, we decided to apply a more stringent cut-off, excluding the 5 Mb regions flanking centromeres for all chromosomes. Applying this new threshold led to an average 35% reduction in the total number of ECSs identified, yet the numbers remained consistent across both reference genomes, with a shift toward the forward orientation (Fig. S4F). Importantly, after this adjustment, binomial tests yielded p-values > 0.05 in both CHM13v2 and RPE-1v1.1 genomes, indicating no significant deviation from random orientation (Supp. Table 6). The refined analysis showed that in both reference genomes, ∼85% of chromosomes now exhibited non-significant deviation from random orientation, highlighting the impact of pericentromeric satellites (Fig. S4G, Supp. Table 7). Indeed, we noted that CENP-B boxes within the centromeric domain have a preferred orientation, usually the opposite to that observed in the flanking pericentromeric, but without the alternance observed along the chromosomes arms (Figures 1 and 2). The ∼15% ECSs that exhibited local orientation biases likely reflect the presence of extended αSat arrays or other islands of repetitive genomic features found throughout the genome that may contribute to these local asymmetries (Fig. S4G, Supp. Table 7). Together, these findings indicate that ECS orientation is globally balanced but locally heterogeneous, with chromosome-specific biases especially within the centromere and pericentromere, but more rarely outside.

### 1.4 ECSs display a non-random spacing pattern conserved across reference genomes

Because ECSs display specific orientation patterns with localized heterogeneity across chromosomes, we reasoned that the boxes may serve as self-hybridizing features for structural organization and/or folding of chromosomes. To further understand whether the ECSs hold functional role(s) within the genome, we analyzed their genome-wide distribution by measuring the linear distance between consecutive occurrences, considering both the same and the opposite orientation, and restricting the analysis to ECSs located more than 5 Mb from centromeres to minimize pericentromeric bias. Genome-wide analysis revealed bimodal spacing distribution, conserved across CHM13v2, and both RPE1v1.1 haplotypes, characterized by a broad range of distances between 0.1-10 megabases (Mb) and a short-range cluster at ∼0.1-3 kilobases (kb) (Fig. 2F), most evident for consecutive ECS pairs with the same orientation. A bimodal pattern for both range of distances was seen only in chromosomes 2 and 9, likely due to a higher number of short-range ECS pairs (Fig. S5, S6). To assess whether this spacing pattern is randomly found, we compared the ECS distances with those of scrambled and αConsensus sequences. For ECSs, long-range spacing (0.1–10 Mb) accounted for more than 70% of all intervals, whereas only a small fraction (∼9%) fell within the short-range cluster at ∼0.1–3 kb (Fig. 2G). In contrast, scrambled controls showed no bimodal pattern (Fig. S7A-C), instead exhibiting a single, narrower spacing distribution spanning 0.1–1 Mb and lacking the short-range cluster characteristic of ECSs. Consistent with this qualitative difference, two–sample Kolmogorov–Smirnov (KS) tests [50,51] confirmed that ECS spacing distributions were significantly different from all controls (p < 0.0001; Supp. Table 8). The αConsensus motif, on the other hand, displayed two separate clusters of distance values: a short cluster between 1–10 bp, corresponding to tightly spaced elements within arrays, and a second cluster between 0.1–1 kb (Fig. S7D-F). The analysis of the short-range group of ECS distances showed an enrichment at 0.1–1 kb (Fig. S4I). This group partially overlaps the second αConsensus cluster, corroborating our aforementioned hypothesis for this cluster, that these ECSs are likely embedded within divergent ectocentromeric αSat arrays on chromosome arms (Fig. S6D-F). Analyzing the long-range group, instead, we observed a significant enrichment at distances spanning 0.1–1 Mb (Fig. S4J). Despite this distribution, the overall ECS spacing remains statistically distinct from αConsensus motif (p-value < 0.0001), as confirmed by KS testing [50,51], indicating that ECS spacing is non-random and not solely determined by αSat arrays (Supp. Table 8). Analysis of individual spacing clusters demonstrated that the short-range cluster showed a more restricted chromosomal distribution. Notably, chromosomes 9 and 2 included the highest enrichment of these short-range distances, partially associated with divergent repetitive elements located beyond 5 Mb from the centromere (Fig. S8A). In contrast, the long-range spacing occurred on all chromosomes, and their relative abundance varied substantially among them (Fig. S8B), consistent with the chromosome-specific heterogeneity previously described for ECS motif orientation. Altogether, these findings demonstrate that ECSs exhibit a statistically significant, non-random spacing pattern.

### 1.5 Motif integrity is associated with ECS conservation across the human population

To investigate the sequence determinants underlying these patterns of ECSs conservation and variability, we next characterized the composition and integrity of the CENNP-B box motif. Sequence logo analysis confirmed a well-defined consensus motif across ECSs positions, consistent with the canonical CENP-B box sequence (Fig. 2H). The motif displays strong conservation across the nine fixed positions, indicating that sequences associated with ECSs follow a highly constrained sequence pattern genome-wide.

We next quantified motif integrity for each ECSs using a Motif Integrity Score (MIS), defined as the number of matches to the nine conserved fixed positions of the CENP-B box motif (Fig. 2I). MIS values therefore provide a quantitative measure of motif fidelity, ranging from fully conserved sequences (score = 9) to highly degenerated motifs (lower scores). Notably, motif integrity differed significantly across ECS conservation categories (Kruskal–Wallis test, p < 2 × 10⁻¹⁶; Fig. 2I). ECSs conserved in most genomes (≥75%) exhibited near-perfect sequence integrity, with median MIS values approaching the maximum score. In contrast, ECSs classified as polymorphic showed intermediate MIS values, indicating partial motif degeneration. Importantly, ECSs absent across all genomes displayed markedly reduced scores, with the lowest values observed for sites located outside alpha-satellite regions (Fig. 2I). These results reveal a strong association between motif integrity and ECSs conservation, indicating that sequence fidelity of the CENP-B box is a major determinant of ECSs stability across individuals. Within alpha-satellite regions, ECSs variability likely reflects structural polymorphism of HOR arrays, whereas outside these regions, reduced motif integrity suggests that a substantial fraction of ECSs represent degenerated or spurious CENP-B box matches. Together, these findings demonstrate that ECSs conservation is not random but is tightly linked to the integrity of the underlying sequence motif, supporting a model in which high-fidelity CENP-B box sequences are preferentially maintained across human genomes.

### 1.6 Intrinsic structural properties of ECS sequences

Our recent work point at centromere sequences forming extremely complex and stable secondary structures, more than other repetitive regions flaking the centromeres and other we analysed [52]. We next investigated whether ECS-associated sequences also exhibit intrinsic structural properties. Sequences flanking ECS motifs display a strong intrinsic propensity to form stable secondary structures. The median minimum free energy (MFE) is −169.4 kcal/mol in haplotype 1 and −168.1 kcal/mol in haplotype 2 (Fig. 2J), substantially lower than shuffled sequences with preserved (−157.1 and −155.5 kcal/mol; p = 4.3 × 10⁻⁸ and 8.0 × 10⁻⁸), indicating increased thermodynamic stability. Consistent results were obtained across three independent scrambled controls based on permuted consensus motifs previously established, all showing significantly higher MFE values (p up to 2.5 × 10⁻¹⁰⁸ in haplotype 1 and 1.7 × 10⁻⁹⁶ in haplotype 2), demonstrating that the observed effect is robust to motif definition and sequence composition. Importantly, this structural signal is highly consistent across the two independent haplotypes, indicating that it represents an intrinsic property of ECS sequences rather than a genome-specific artifact. Stratification of ECS pairs by relative motif orientation (Same vs Opposite) revealed no significant differences in either haplotype (haplotype 1: p = 0.971; haplotype 2: p = 0.826), excluding a contribution of inter-motif interactions such as inverted repeat pairing (Fig. S4H). Together, these results demonstrate that the propensity for secondary structure formation is an inherent feature of the local ECSs context. Collectively, these findings provide a mechanistic basis linking the non-random genomic distribution of ECSs to intrinsic sequence-encoded structural properties, supporting the hypothesis that ECSs may contribute to genome organization through their biophysical potential.

### 1.7 ECS occupy specific positions within three-dimensional (3D) genome architecture

We next examined whether the intrinsic structural propensity of ECSs is associated with specific features of three-dimensional genome organization. To this end, we analyzed haplotype-resolved Hi-C data to characterize the chromatin context of ECS loci. We performed isogenomic conformation mapping [38] using the RPE-1v1.1 reference to align data from a publicly available Hi-C experiment generated in RPE-1 cells [14]. Chromatin compartments and topologically associating domains (TADs) were identified at 20 kb resolution for both haplotypes.

Visual inspection of Hi-C contact maps revealed that ECSs are distributed across different chromatin contexts, including intra-TAD regions, TAD boundaries, and boundary-proximal sites in both haplotypes (Fig. 3A). This qualitative observation suggests that ECS are not confined to a single structural compartment but may preferentially localize to regions involved in chromatin domain organization. To further test whether this localization is non-random, we performed a permutation analysis preserving chromosomal distribution (Fig. 3B). ECS were found significantly closer to TAD boundaries than expected by chance (median distance 188.8 kb and 186.1 kb versus 210.0 kb and 208.2 kb in randomized controls; empirical p < 0.001 and < 0.002). This independent approach confirms that ECSs preferentially occupy genomic regions in proximity to chromatin domain boundaries. In addition, ECSs are systematically located at genomic positions with features of local chromatin domain separation (Fig. 3C). The median insulation score is −4.20 in haplotype 1 and −4.18 in haplotype 2, significantly more negative than all scrambled controls (p up to 4.8 × 10⁻¹⁰⁰ and 4.6 × 10⁻¹⁰⁵, respectively), indicating reduced local chromatin interconnectivity at ECSs positions. This pattern is consistent across both haplotypes and independent of motif definition. Together, these results demonstrate that ECSs are not only characterized by intrinsic structural properties encoded in their sequence, but are also positioned at specific sites within the three-dimensional genome architecture, corresponding to regions of domain separation. This convergence between sequence properties and spatial genome organization strongly supports a structural role for ECSs in genome organization, despite the absence of stable loop formation detectable at bulk Hi-C resolution. Their position often with alternating CENP-B boxes in inverted orientation on the same strand may act analogously to Alu inverted repeats, promoting self-complementary base-pairing, chromatin looping and topological alterations that influence local gene regulation.

**Figure 3.**
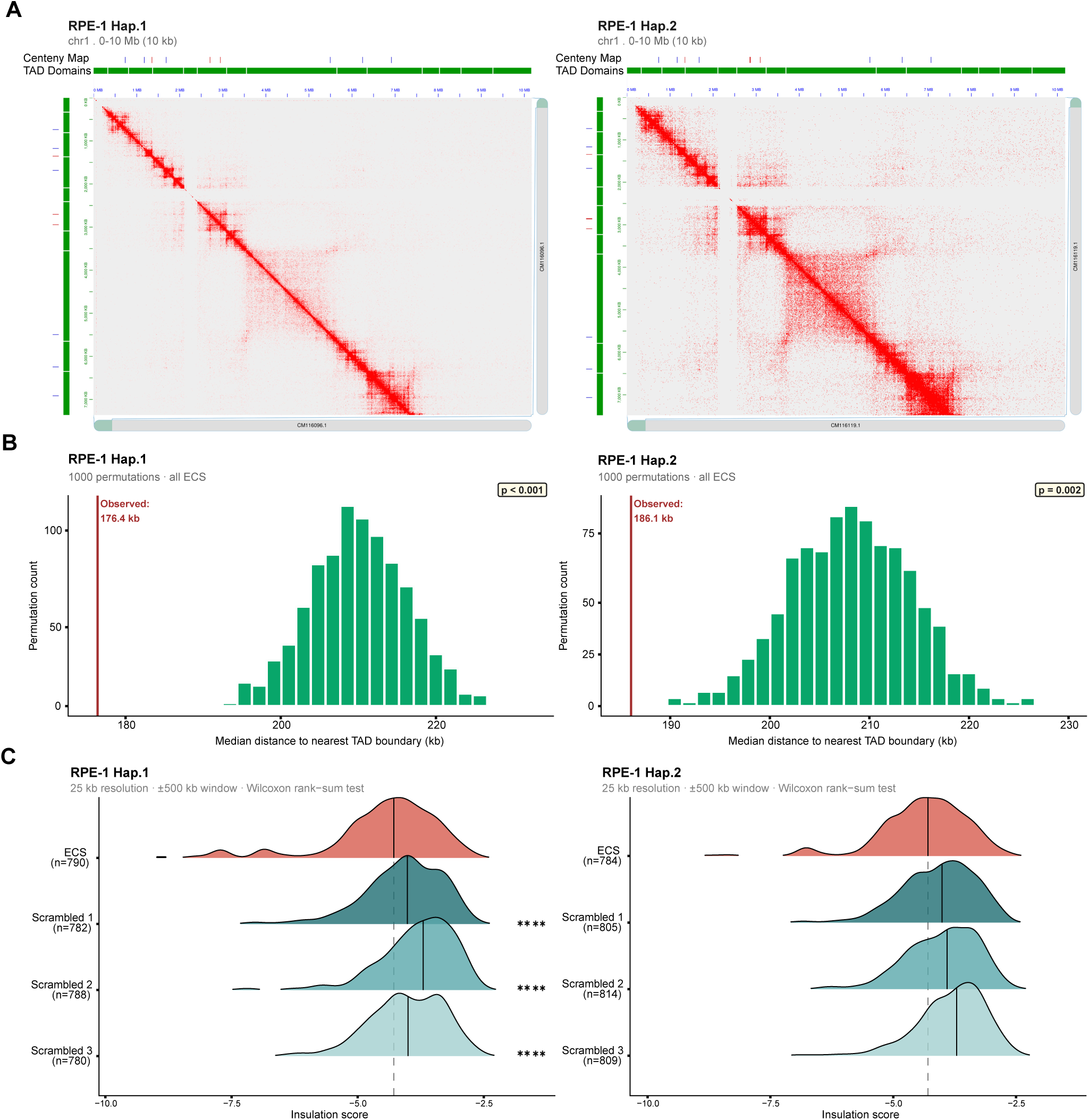
The 3D organization of ECS loci shows non-random positioning near chromatin domain boundaries. A) Representative Hi-C contact maps generated from RPE-1v1.1 haplotype 1 (A) and haplotype 2 for chromosome 1, visualized using Juicebox. The contact enrichment signal is shown as a heatmap. TAD domains are displayed as green rectangles on the left and top tracks, together with the human centeny map showing ECS motifs in forward (blue) and reverse-complement (red) orientation. No preferential enrichment of Hi-C contacts is observed at consecutive inverted ECS pairs relative to surrounding chromatin. B) Permutation test of ECS distance to TAD boundaries. Histograms show the distribution of permuted median distances (1000 iterations) from ECS loci to the nearest TAD boundary, for haplotype 1 (left) and haplotype 2 (right). Vertical line marks the observed median for real ECS loci, significantly closer than expected (188.8 / 186.1 kb observed vs 210.0 / 208.2 kb permuted; p < 0.001 / < 0.002). C) Insulation score at ECS loci. Ridgeline plots showing the distribution of insulation scores (25 kb resolution, log₂(cross / √(left × right))) at ECS loci versus three scrambled control sets, for haplotype 1 (left) and haplotype 2 (right). ECS loci show significantly more negative scores than all controls in both haplotypes (median −4.20 / −4.18; two-sided Wilcoxon p up to 4.8×10⁻¹⁰⁰ / 4.6×10⁻¹⁰⁵).

### 1.8 Occurrence of ECSs within gene features and chromatin accessibility

To address ECSs functional relevance, we first investigated their localization within genomic features, including gene-associated regions. We had previously found ∼40% of ECSs located within gene features, with ∼9% associated with promoter regions, representing moonlighting function of CENP-B outside of centromeres in transcription [20]. Here, we further examined whether ECSs associate with transcriptionally accessible chromatin by intersecting their genomic coordinates with publicly available ATAC-seq data from hTERT-RPE-1 cells [28]. Analysis across both RPE-1 haplotypes showed that only ∼5% (48 in haplotype 1 and 49 in haplotype 2) colocalize with ATAC-seq peaks (Fig. S11A). Among the accessible subset, ∼20% are located in intergenic regions, ∼40% in promoter regions and, the remainder distributed across exons, introns, and transcription start sites (TSSs) (Fig. S11A). These results indicate that ECSs are predominantly found in inaccessible chromatin, but a small subset is enriched in gene-proximal and transcriptionally active regions. This pattern supports a model of functional heterogeneity, in which a subset of ECSs may retain context-dependent regulatory potential, potentially enabling protein recruitment in permissive chromatin environments. This observation prompted us to further investigate whether such heterogeneity reflects differential CENP-B binding and chromatin organization, especially in light of a recurrent state of a region of chromatin accessibility preceded by enrichment in trimethylated lysine 27 in histone H3 (K27me3). The organization of the chromatin states underlying and neighboring the ECS is also intriguing. Compacted marks K27me3 precedes the region of chromatin accessibility that is flanked by CENP-B boxes and perfectly overlaps with CENP-B peaks, often multiple ones. The peaks do not seem to perfectly localize with the motifs, another interesting observation, suggesting that the conformation of chromatin is organized three-dimensionally more than how we visualize it in a linear fashion from *p* to *q* arm.

### 2.1 From catalog expansion to functional classification of “ectocentromeric sites” based on CENP-B binding and chromatin context

Having established that ECSs are highly conserved across individuals, under negative selection, and exhibit non-random spatial organization despite lacking stable inter-ECS genomic interactions, we next investigated whether this evolutionary constraint reflects a latent functional capacity related to centromeric protein recruitment. Specifically, we asked whether a subset of ECSs retains the ability to bind CENP-B outside the centromere, and whether their chromatin state influences such binding. To address this, we integrated CENP-B CUT&RUN profiles from hTERT-RPE-1 cells [53] with ECS coordinates across both RPE-1v1.1 haplotypes and ATAC-seq accessibility data [28], enabling a functional classification of ECSs based on protein occupancy and local chromatin state (Fig. 4A; Fig. S9; Fig. S10). This analysis allowed us to classify these CENP-B box elements into different categories, here renamed “*ectocentromeric sites*”, to denote loci functionally defined by their association with CENP-B motif enrichment, CENP-B box position, and chromatin state (Fig. 4A-B). This classification includes: (i) *box-only*, where only the motif is present. This category accounts for approximately 84% when considering only the box subset, but only 42% when all four categories are considered; (ii) *box-protein*, referring to sites directly associated with CENP-B. This category is relatively infrequent, accounting for approximately 6% of all ectocentromeric sites identified, and is mostly found within divergent αSat arrays located either in pericentromeric regions or beyond 5 Mb from the centromere; (iii) *box-near-protein*, which includes sites located mainly in a range of 0.1-1 Mb away from the nearest CENP-B peak, typically associated with open chromatin (Fig.S11B). This category accounts for ∼1% of all ectocentromeric sites. Notably, this distance range overlaps with the most prevalent long-range spacing observed between consecutive ECSs (0.1-1 Mb), suggesting that local ECS architecture could influence CENP-B positioning; (iv) *protein-only* sites, comprising ∼700 significant CENP-B peaks identified outside centromeres, which occur at loci lacking the canonical CENP-B box motif. This category represents roughly 50% of all sites when considering all four categories across both haplotypes. These non-canonical sites span diverse genomic features and chromatin states, representing atypical binding events. Overall, this classification establishes that ectocentromeric sites exhibit substantial heterogeneity in their protein occupancy and chromatin state, ranging from structurally silent *box-only* elements to *protein-only* sites actively bound by CENP-B of which ∼30% are in accessible chromatin. While chromatin accessibility reflects the local regulatory environment at each site, we next asked whether these categories differ in their chromatin context, we next examined their distribution across A/B compartments derived from Hi-C data. Each ECS was assigned to either compartment A (active, open chromatin) or compartment B (repressed chromatin), and the fraction of sites in compartment A was compared across categories (Fig 4C). Strikingly, the four categories displayed a progressive enrichment in compartment A that correlates with CENP-B binding stability. *Box-only* and *box–near–protein* sites showed an approximately random A/B distribution (∼50% in A), indicating that the presence of the motif alone does not influence compartment localization. In contrast, *protein-only* sites exhibited a significant enrichment in compartment A (∼57–65%), while *box–protein* sites showed the strongest enrichment (up to ∼94% in A), despite their low abundance. Together, these results reveal a clear dose–response relationship in which increasing stability and specificity of CENP-B binding correspond to a stronger association with open chromatin compartments, suggesting that stable CENP-B occupancy is favored in, or constrained by, accessible chromatin environments.

**Figure 4.**
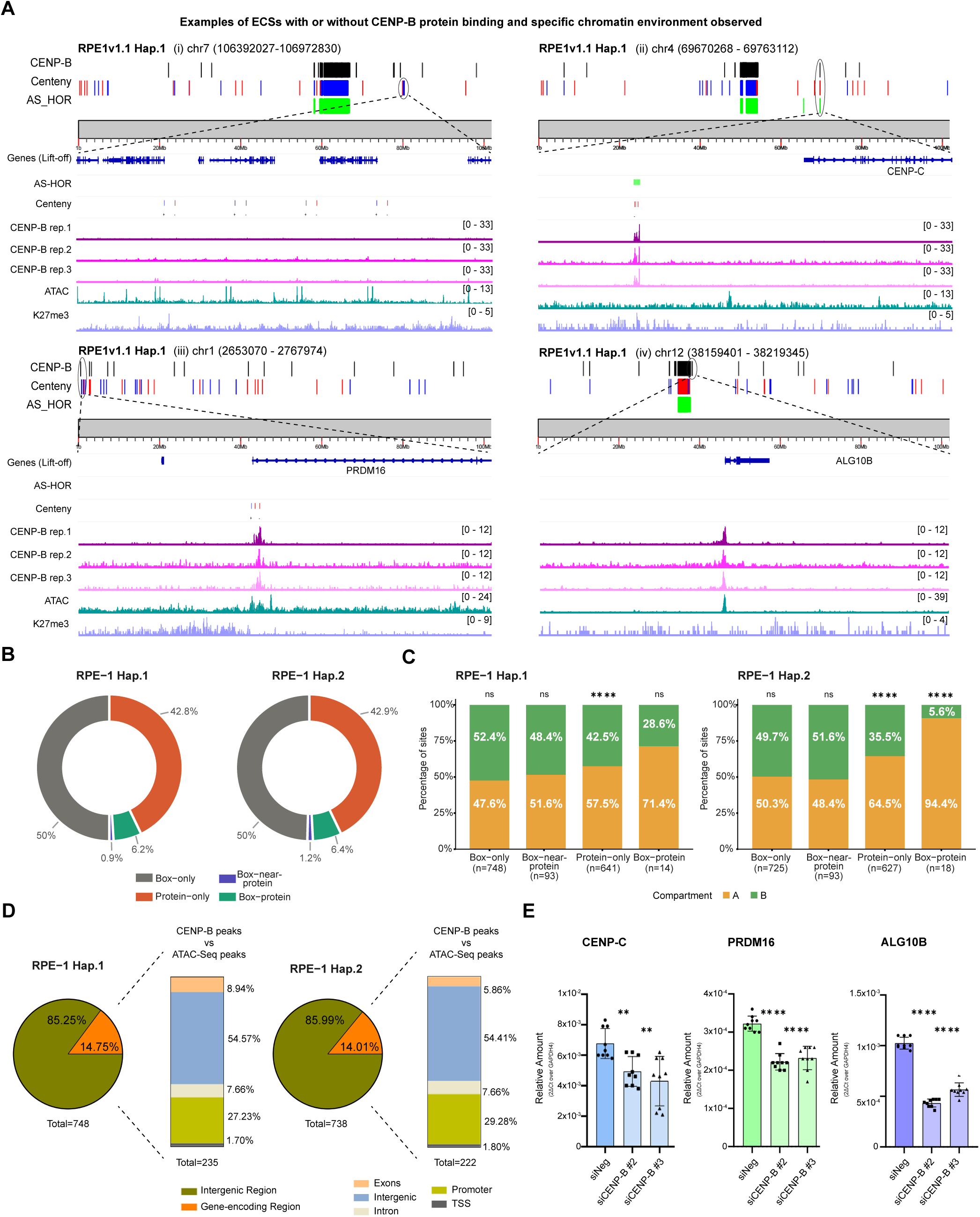
Ectocentromeric sites classification including CENP-B binding outside the centromere. A) Representative IGV views and ideograms of ecotentromeric sites corresponding to four categories: (i) box-only, (ii) box–protein, (iii) box-near-protein, and (iv) protein-only sites, showing CENP-B CUT&RUN peaks and α-satellite annotation. Ideograms depict the genomic context of each box category together with CENP-B peaks and α-satellite tracks, while IGV snapshots provide locus-level detail. Gene annotations (RPE-1v1.1) were transferred from ENSEMBL hg38 via liftOver. CENP-B CUT&RUN peaks were treated as broad regions, and signal tracks were normalized to input controls. B) Donut charts represent the percentage distribution of ectocentromeric sites across the four categories. C) Barplot showing the percentage of ectocentromeric sites in compartment A for each functional category (box-only, protein-only, box-protein, box-near-protein)), for both haplotypes. D) Pie charts summarizing the percentage of ECSs found in intergenic or coding regions, and bar plots indicating the CENP-B peaks distribution across genomic features and colocalized with ATAC-Seq signal. E) qPCR analysis of selected genes positioned on the previously described chromosomes following CENP-B knockdown (** p-value < 0.005, **** p-value < 0.0001).

### 2.2 CENP-B occupancy outside the centromere reveals moonlighting function in transcriptional regulation

Our previous analysis of the CUT&RUN of CENP-B [53], K27me3 [29], and ATAC-seq datasets [28], found that the CENP-B read density displays a clear bimodal distribution on either side of the transcription start sites (TSSs), highlighting the potential role in transcriptional gene regulation [20]. Building on these observations, we quantified the extent of CENP-B binding outside centromeres. We identified 748 significant peaks in haplotype 1 and 738 peaks in haplotype 2 located outside the centromere (Fig. 4D). Notably, ∼85% of significant peaks were in intergenic regions in both haplotypes, while approximately 15% overlapped coding regions. Moreover, ∼30% of these significant peaks described above colocalize with ATAC-seq signals, mainly located within intergenic (∼54%) and promoters (∼30%) regions. Only ∼2% of occurrences represent CENP-B binding at the TSSs (Fig. 4D), with CENP-B bimodal distribution around promoters thus limited to a subset of sites. In addition, some ECSs located in proximity to the TSSs of specific genes, ranging from ∼50 kb to ∼4 Mb away (Supp. Table 9). These include genes encoding zinc-finger proteins, such as *ZNF716*, which is involved in transcriptional regulation [54]; *TRIM51G*, a member of the TRIM family, which participates in ubiquitination, DNA repair, and genome surveillance, with some TRIM proteins being associated with chromosomal rearrangements in cancers [55]; *CENP-C*, which is a constitutive component of the Constitutive Centromere-Associated Network (CCAN) that directly binds CENP-A nucleosomes and bridges the inner centromere with the outer kinetochore, playing an essential role in chromosome segregation [56,57]. Through its direct interaction with CENP-B, CENP-C contributes to centromere integrity and kinetochore stability [17]; and *OR11H1*, an olfactory receptor gene that belongs to a large, repetitive gene family. Many of these loci are embedded within segmental duplications, making them hotspots for genomic rearrangements and structural instability [58]. The recruitment of CENP-B in proximity to the regulatory regions for these genes makes this findings even more interesting. To explore the potential functional consequences of CENP-B binding at ECSs, focusing on possible involvement in gene regulation, we selected three representative CENP-B-bound sites from full set identified by CUT&RUN [53] (Supp. Table 10, 11). These include CENP-B bound to canonical ECSs embedded within αSat array, ∼70 kb away from the TSS of the *CENP-C* gene (ii); CENP-B bound to canonical ECSs located within the first intron of the *PRDM16* gene and colocalized with significant ATAC-Seq peaks. Since these ECSs are closely spaced, this suggests that CENP-B may bind all nearby ECSs (iii); CENP-B bound to non-canonical ectocentromeric site on the *ALG10B* gene promoter, located within an accessible chromatin region (iv). We then knocked down CENP-B using two previously validated siRNAs (Fig. S11C), followed by quantitative PCR (qPCR) to measure the relative expression of the previously chosen genes described above. The qPCR analysis revealed a significant downregulation of all three genes upon CENP-B knockdown compared with the negative control (Fig. 4E). ALG10B showed the strongest reduction (∼2-fold, p < 0.0001), followed by PRDM16 (∼1.4-fold, p < 0.0001) and CENP-C (∼1.3-fold, p < 0.002), all normalized to GAPDH. Although the full mechanistic details remain to be elucidated, the differential downregulation observed across PRDM16, CENP-C, and ALG10B genes upon CENP-B depletion suggests that CENP-B influences transcription through distinct mechanisms depending on the local chromatin context, acting directly at canonical ECS loci and indirectly at non-canonical binding sites through interactions with other regulatory factors.

### 2.3 CENP-B binding state defines local 3D contact patterns between ectocentromeric sites

To investigate whether ECSs interact in three-dimensional space, we analyzed Hi-C contact frequencies between pairs of consecutive *box-only* sites and *box-near-protein* sites using Aggregate Peak Analysis (APA). Strikingly, distinct interaction patterns emerged depending on the CENP-B binding status of the sites. *Box-only* sites exhibited a strong depletion of Hi-C contacts at short genomic distances (0–100 kb) (Fig. 5A), with a pronounced reduction compared to distance-matched random pairs (fold-change <1, p < 10⁻¹⁴, large negative effect size) (Fig. 5C). In contrast, *box–near–protein* pairs showed significant enrichment of contacts within the 100–200 kb range (fold-change >1, p < 0.01, large positive effect size) (Fig. 5D), indicating enhanced spatial proximity when CENP-B is bound to one of the elements. These opposing patterns define a clear double dissociation, whereby the absence of CENP-B binding is associated with local contact depletion, whereas its presence promotes focal enrichment of chromatin interactions. Importantly, both effects were independently replicated in each haplotype, with consistent fold-change direction, effect size, and statistical significance, demonstrating that these patterns are not driven by haplotype-specific genomic configurations (Fig. S12A-B). Extending the analysis across the full 0–1 Mb range revealed that these effects are strongly distance-dependent (Fig. S12C). *Box-only* sites showed progressive contact depletion extending up to ∼400 kb, whereas *box–near–protein* sites displayed enrichment restricted to ∼200 kb, beyond which interactions returned to baseline levels. Together, these findings uncover a previously unrecognized layer of ECSs organization in which the presence of CENP-B binding may shape local chromatin folding. We propose that stable or partial CENP-B occupancy promotes the formation of specific short-range contacts, whereas its absence results in a more dispersed or repulsive spatial configuration, highlighting a structural role for CENP-B beyond its canonical centromeric function.

**Figure 5.**
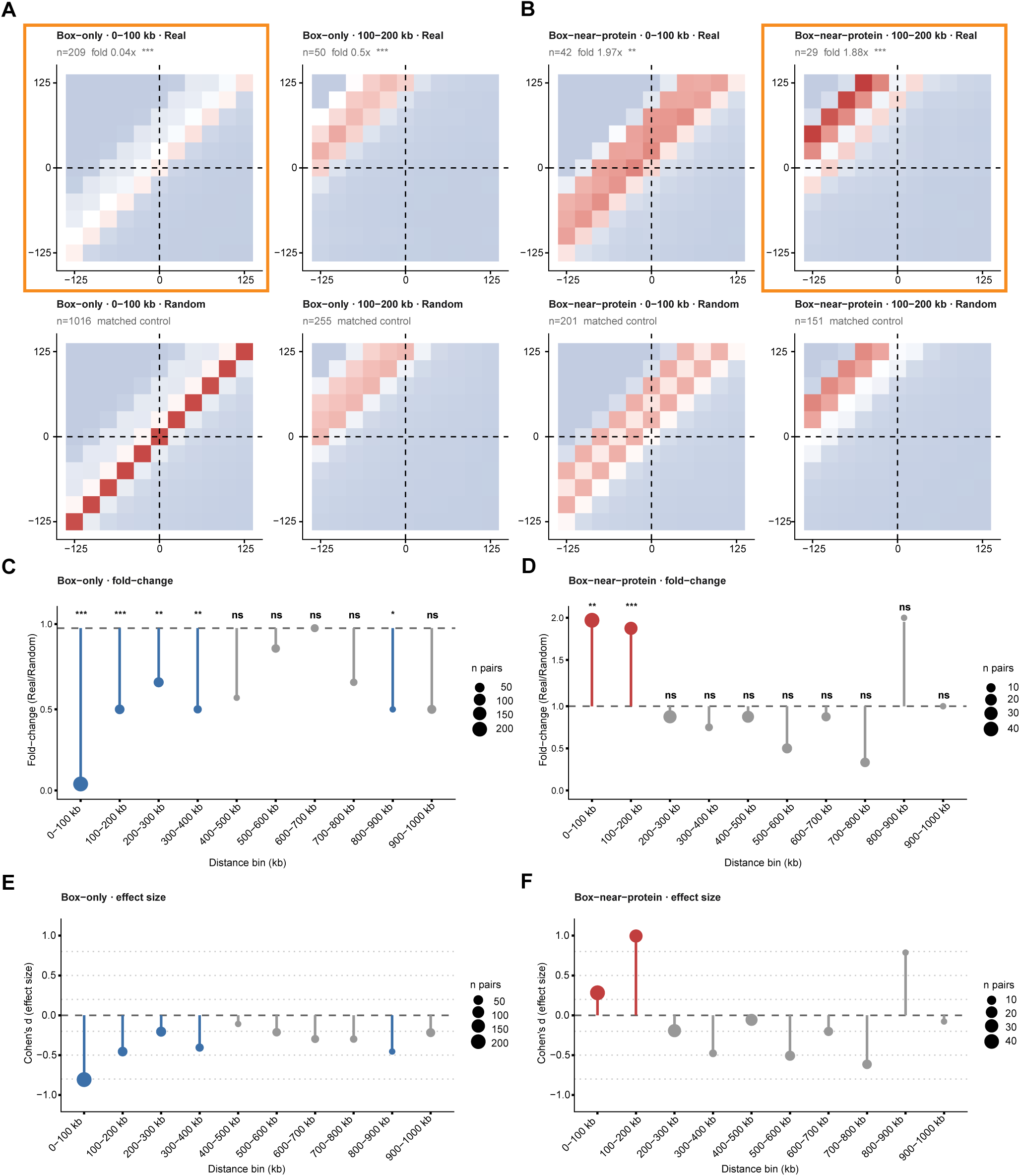
Distance-stratified APA reveals a double dissociation between Box-only and Box-near-protein pairs. A-B) APA heatmaps (11×11 bin, ±125 kb, 25 kb resolution) of Box-only (BO) (A) and Box-near-protein (BNP) (B) pairs at the two key distance bins (0–100 and 100–200 kb), comparing real (top) and distance-matched random (bottom) pairs. Orange-bordered panels mark the bin of maximum effect per category. (C-D) Lollipop plots showing fold-change (real/random of central [6,6] values) across 10 distance bins for Box-only (C) and Box-near-protein (D); point size scales with pair number. (E-F) Lollipop plots showing Cohen’s d effect size across distance bins for Box-only (E) and Box-near-protein (F). Wilcoxon rank-sum test (two-sided for Box-only, one-sided for Box-near-protein); ***p < 0.001, **p < 0.01, *p < 0.05, ns p ≥ 0.05.

### 2.4 Relationship between ectocentromeric sites and human neocentromeres

Transcription has been associated with both seeding of neocentromere [59] and for endogenous centromere function [60,61]. To this end, we characterized ECSs in relation to neocentromeric loci using all previously mapped sites. Visualization of publicly available CUT&RUN tracks for CENP-A, H3K9me3, and H3K27me3 [62], overlaid with ECS coordinates, revealed the consistent presence of the ECSs cluster several megabases away from the experimentally induced Neo4p13 site [62,63]. Specifically, an ECS cluster spanning 1-3 Mb from the CENP-A domain was detected in early passage cells (day=0). Upon SUV39H1/H2 and SUZ12 inactivation, an independently maintained cell population proliferated for 100 days exhibited a reduction in this distance, caused by neocentromere drift and expansion of ∼150 kb into adjacent permissive chromatin (Fig. 6A). Although ECSs are likely too distal to directly influence neocentromere formation, their spatial proximity to neocentromeric boundaries, together with the time-dependent expansion of CENP-A domain, suggests that ECSs may act as latent secondary boundaries that contribute to long-term neocentromere stabilization. Similar ECS arrangements were observed at recently characterized clinical neocentromere, including those at 4q21.3, 8q21.2, and 13q21.1 (Fig. 6B) [64]. Notably, the bilateral enrichment of the ECS cluster at 8q21.2 further suggests that this mechanism may operate symmetrically, constraining neocentromere expansion in both directions. These insights raise the possibility that ECSs may acquire functional relevance in genomic contexts characterized by intense chromatin remodeling, thereby extending their biological impact well beyond their basal structural and regulatory roles.

**Figure 6.**
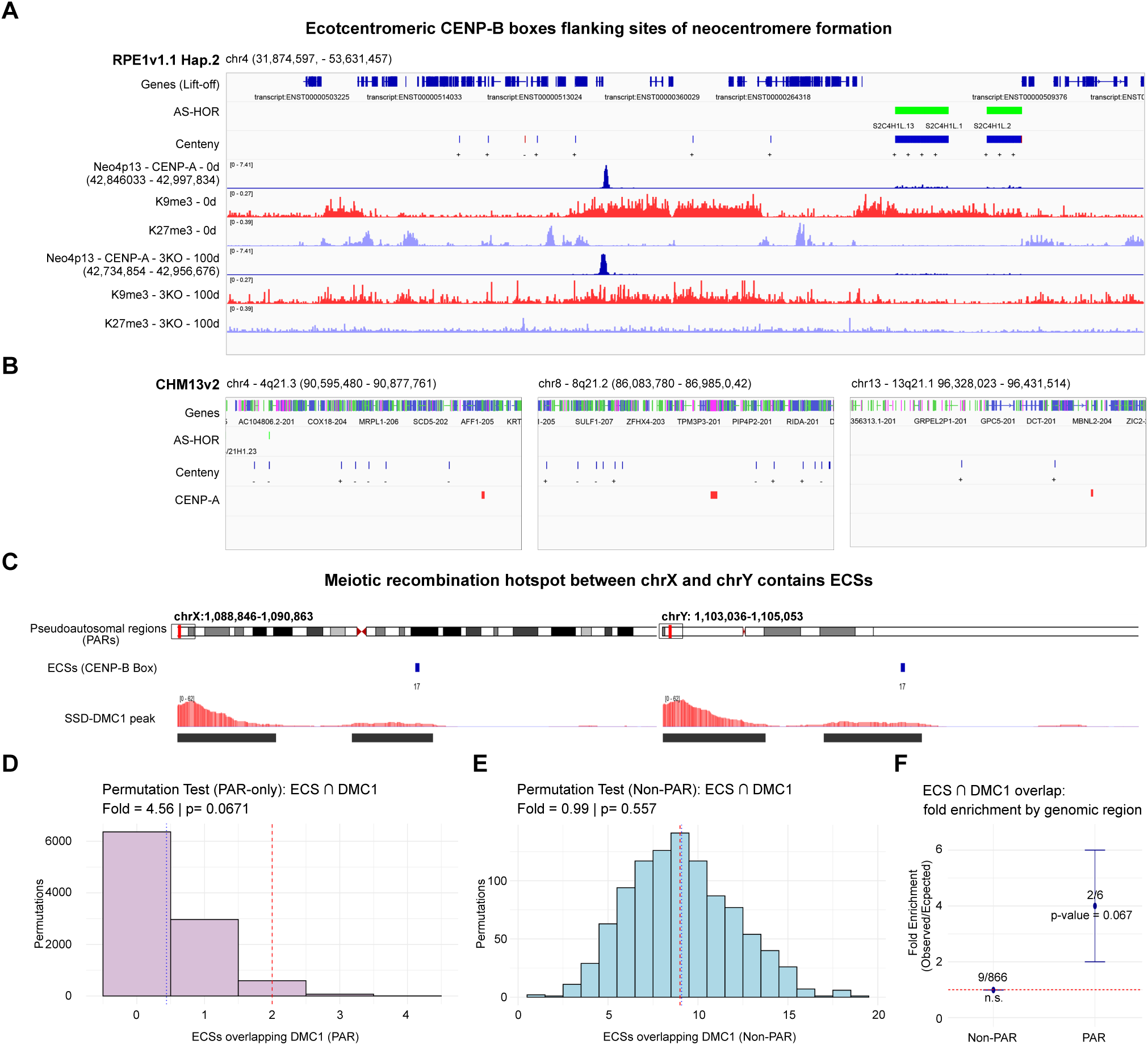
Context-dependent functional associations of ECSs at neocentromere boundaries and meiotic recombination sites’. A–B) Representative IGV snapshots illustrating the spatial relationship between selected neocentromeric sites, including (A) the experimentally induced Neo4p13 mapped in RPE1v1.1 haplotype 2 and (B) the clinical neocentromeres at 4q21.3, 8q21.2, and 13q21.1 identified in CHM13v2. In Neo4p13, tracks show colocalization with enrichment for H3K9me3 and H3K27me3. All neocentromeric regions, defined by CENP-A peaks, are displayed across multiple time points (0–100 d) and with or without (3KO) SUV39H1/H2 and SUZ12 expression. C) Representative images showing overlaps between ECSs and DMC1 SSD-Seq peaks within the PARs of chromosomes X and Y. Tracks display ECS coordinates and DMC1 enrichment signals within PAR1 and PAR2. D) Permutation analysis restricted to PAR coordinates. The histogram shows the distribution of overlaps obtained after 10,000 random repositioning of ECS intervals, while preserving chromosome identity and element size. The blue line indicates the mean expected overlap, whereas the red line marks the observed value. The observed count falls in the upper tail of the null distribution, corresponding to an approximately fourfold enrichment (z = 2.09; empirical p = 0.067). E) Permutation analysis outside the PAR. Using the same framework but restricting randomization to non-PAR genomic space, the observed number of overlaps lies near the center of the null distribution (fold ≈ 0.99; z = –0.03; empirical p = 0.557), indicating no enrichment. F) Summary dot plot of fold enrichment (observed/expected) across genomic contexts. The dashed line indicates the null expectation (fold = 1). ECSs show preferential enrichment within PARs, whereas no deviation from expectation is detected genome-wide outside these regions.

### 2.5 DMC1-marked recombination sites preferentially associated with ECSs in the pseudoautosomal regions

One of the most interesting feature of ECS is their conserved position. Notably, these overlaps occurred within the pseudoautosomal regions (PARs) on chromosome X and Y, where meiotic recombination is obligatory and particularly frequent (Fig. 6C), where the first 3 CENP-B boxes starting from the sex chromosomes’ *p*-arm are perfectly conserved in sequence [20]. This is particularly interesting considering that endogenous centromeres are considered refractory to recombination. To explore the possibility that these ECS may be involved in meiotic recombination, we compared the spatial association between ECSs and DMC1, a recombinase that marks active meiotic break sites and defines recombination hotspots genome-wide. Analysis of publicity available DMC1 SSD-Seq dataset [37] identified 11 overlaps between ECSs and DMC1 peaks that occur in multiple chromosomal contexts. A permutation analysis that preserves chromosome identity and interval size demonstrated that this association is not uniformly distributed across the genome. Specifically, within the PAR regions, 2 out of 6 ECSs overlapped DMC1 peaks. When ECSs were randomly repositioned within the PAR in 10,000 permutations, the resulting overlaps were consistently lower than the observed value, indicating a ∼4-fold enrichment (z = 2.09; p = 0.0671) (Fig. 6D). This supports a trend toward preferential association between ECSs and DMC1 in the PAR. However, given the small number of ECSs within the PAR (n=6) and the resulting limited statistical power, this finding remains preliminary and requires validation in larger datasets. In contrast, outside the PAR, only 9 overlaps were observed among 1359 ECSs identified in the CHM13. Under the same permutation framework restricted to non-PAR genomic space, the observed value did not deviate from random expectation (fold change ≈ 0.99; z = -0.03; p = 0.557) (Fig. 6E). The absence of enrichment outside the PAR indicates that association with DMC1 is not a general property of ECSs across the genome (Fig. 6F).

Together, these analysis supports a context-dependent model of ECS function, in which the majority of sites remain structurally silent under basal conditions, while specific subsets, defined by CENP-B occupancy, chromatin accessibility, or exposed to intense chromatin remodeling, exhibit distinct functional properties ranging from gene regulation to potential roles in meiotic recombination.

## Discussion

This study provides the first comprehensive characterization of ectocentromeric sequences and expands the classification beyond just CENP-B boxes to ectocentromeric sites including occurrences of CENP-B protein enrichment in absence of the underlying motif seen in chromatin-immunoprecipitation peaks along chromosome arms. Ectocentromeric sites (ECS) represent functionally heterogeneous genomic elements, establishing that their evolutionary conservation reflects not a single function, but a broad range of activities influenced by the chromatin context, spanning from higher-order chromatin organization and gene regulation to potential roles in meiotic recombination. Although most ECSs are epigenetically silent under basal conditions, their conserved genomic positions, orientations, and spacing patterns indicate that this organization is unlikely to be functionally irrelevant. Building on previous work on the characterization of the CENP-B box outside the centromere [20], our findings redefine these elements as modulators of chromosome architecture, whose activity becomes evident only under specific chromatin states and cellular conditions. For our identification of the motif on chromosome arms, we chose a degenerate version (5’ NTTCGNNNNANNCGGGN 3’), which contains 8 degenerate and the 9 fixed nucleotides known to be important for the physical docking of CENP-B. The CENP-B box motif (5’ YTTCGNNNNANRCGGGN 3’) is known to represent a high-affinity binding site for CENP-B [24,65]. When applying the canonical CENP-B binding motif, we observed only marginal differences (∼3%) in ECS detection, spacing distributions, and chromosomal patterns compared to our selected ECS motif (Supp. Tables 12-14). These results indicate that the ECS motif used for this study effectively captures the spatial and organizational features of genomic regions associated with CENP-B occupancy, including all experimentally detected ectocentromeric binding sites, while remaining independent of binding affinity considerations. Consistent with this model, we found both ECS that do not bind CENP-B and CENP-B-only sites, where the protein enrichment is not over underlying motif. We also find significant amount of ECS where the motif is adjacent but not directly underlying the CENP-B protein enrichment, indicating that chromatin state and/or conformation more than direct binding to this sequence, dictates interaction between boxes and protein outside of centromeres. Importantly, the presence of these degenerate CENP-B boxes found on chromosome arms outside the centromeres and away from adjacent pericentromeric repeats appears to be under strong evolutionary constraint.

The genomic distribution of CENP-B box motifs outside centromeres provides compelling evidence for negative selection shaping their occurrence. While degenerate motifs with identical nucleotide content to CENP-B boxes are found stochastically across the genome at an estimated 10,000–40,000 loci, ectocentromeric CENP-B boxes are retained at only ∼1,000 positions representing a 10 to 40-fold depletion relative to the sequence-permissive background. This striking deficit indicates that the vast majority of potential ectopic CENP-B binding sites are actively purged from the genome, while a defined subset is maintained at stable, chromosome-specific positions and with specific orientation across individuals. Crucially, the retained sites are not randomly distributed drifting sequences: their positional conservation across complete T2T assemblies, their enrichment at TAD boundaries, meiotic recombination sites, and regions of context-dependent chromatin accessibility, and the functional consequence of their loss such as reduced local gene expression upon CENP-B knockdown for those ECSs where both boxes and protein are found, collectively argue that these loci are preserved by selection because they are useful, not randomly. Together, these data reframe ectocentromeric sites from incidental sequence occurrences to an evolutionarily constrained regulatory layer embedded within the architecture of human chromosomes.

We found that ECSs are largely conserved across multiple human near-T2T chromosome level assemblies, with only a small fraction representing non-canonical sequences compared to the consensus CHM13 *centeny* map. Whereas complete T2T assemblies reveal positional conservation, contig-level assemblies from large cohorts show considerable apparent variation. There are many plausible explanations from consequences of contig fragmentation, assembly errors, ambiguous anchoring of contigs to chromosome coordinates that make recognition with respect to CHM13 as “reference” difficult and/or *bona fide* biological divergence and polymorphism as we extend from just a handful of curated haplotypes to hundreds of genomes. While polymorphism of ECS position and orientation is undoubtably evident, their exquisite conservation in T2T genome spanning diverse ancestry points toward the divergence being modest and most ECS being largely present in the expected orientation within a specific region +/- 1 Mb. Further, polymorphic ECS may be interesting on their own right with functional significance still unclear, including their potential effects on phenotype, fitness, or disease susceptibility. However, their study may potentially uncover novel genetic diversity, provide insight into evolutionary and functional genomics, and be leveraged to investigate evolutionary mechanisms.

Comparative analyses with T2T assemblies from humans and primates further revealed that ECSs predominantly occupy canonical genomic positions in humans, whereas in primates they exhibit chromosome-specific conservation patterns. This suggests selective maintenance of specific loci across evolutionary lineages and points to a potential role in structural genomic organization. Analysis of ECS features revealed non-random orientation and a conserved bimodal spacing pattern across both CHM13 and RPE-1 genomes. Although these properties are globally conserved, the distribution of orientations and the relative abundance of long- and short-range spacing vary across individual chromosomes. This aspect indicates that each chromosome exhibits a unique arrangement of ECSs shaped by local sequence composition and the presence of repetitive elements, which may be critical for understanding their role in chromatin organization. The enrichment of ECSs at 0.1–1 Mb and 0.1–1 kb distances identified likely reflects 3D chromatin organization and suggests a potential role for ECS distribution in shaping discrete structural domains within the genome. Their highly conserved genomic position, orientation bias, and spacing patterns are consistent with a model of probabilistic, context-dependent chromatin modulation, where ECSs may influence local chromatin organization only under specific genomic or cellular conditions, potentially contributing to regulatory mechanisms such as enhancer-promoter contacts and facultative heterochromatin boundaries. This hypothesis was supported by the absence of stable Hi-C contacts at ECS pairs, which distinguishes them from canonical loop anchors such as CTCF-cohesin sites, placing them instead in a growing class of genomic elements whose organizational properties are compatible with rare or conditionally active interactions that fall below the sensitivity of ensemble Hi-C. The integration of CENP-B CUT&RUN and ATAC-seq data allowed us to better resolve this functional diversity by defining distinct categories collectively referred to as “*ectocentromeric sites*” because they include both ectocentromeric sequences (CENP-B boxes) and sites of CENP-B recruitment in absence of underlying DNA motif. The presence of reverse complement CENP-B box sequences on the same DNA strand, together with the observed non-random orientation and spacing in some chromosomes, could promote the formation of secondary structures enriched at centromeres and along chromosome arms [18,66]. These features are consistent with the probabilistic model described above, in which the formation of micro-interactions arise within a specific chromatin context that fall below the detection threshold of ensemble Hi-C. Notably, only a small fraction of the *box-only* category (∼5%) resides within accessible chromatin, as previously described. While the majority of these sites likely contribute to higher-order chromosomal architecture, this accessible subset may be functionally distinct and potentially engaged in regulatory roles, despite the absence of detectable CENP-B binding. Based on the observed CENP-B binding pattern and functional validation through knockdown experiments, we provide evidence supporting CENP-B moonlighting function(s) outside of its primary role within the centromere. Consistent with our findings, recent work has shown that CENP-B can bind non-centromeric sites in chromosome arms, including promoters, independently of the canonical B box motif, through recognition of DNA secondary structures such as hairpins, and can influence gene expression [67]. The differential rate of downregulation observed across PRDM16, CENP-C, and ALG10B genes upon CENP-B knockdown suggests that CENP-B influences gene expression through distinct mechanisms depending on the local chromatin context. This mechanism may include direct binding at canonical ECS loci within gene bodies or promoter-proximal regions, as well as indirectly at non-canonical binding sites mediated through interactions with other regulatory factors. Interestingly, CENP-B has been shown to promote DNA loop formation at centromeres through its dimerization domain [18], raising the possibility that a similar looping mechanism may operate at “*ectocentromeric sites”*, potentially mediating long-range chromatin contacts between ECS-bound CENP-B and distal regulatory elements. The regulation of another centromeric protein, CENPC, is particularly intriguing and may be part of a feedback loop or cell cycle dependent regulation/sequestration of CENP-B away from ectocentromeric sites to its canonical role at centromeres during mitosis. Together, these findings suggest that these newly identified sites outside the centromere may fulfill multiple roles, with most contributing primarily to chromatin structural organization, whereas a distinct subset may modulate gene regulation via CENP-B binding or by modulating local chromatin accessibility.

Beyond chromatin organization and gene regulation, our analyses suggest that ECSs may also acquire additional functional relevance under specific genomic conditions. We can attempt to speculate that drift and expansion of the CENP-A domain within a neocentromeric region toward ECS clusters may enable CENP-B binding, thereby recruiting the DAXX/ATRX complex and promoting H3.3 deposition together with local heterochromatin reinforcement through residual H3K9me3 and HP1 compaction. This process may contribute to restricting further neocentromere expansion when canonical heterochromatin pathways are compromised. Presence of CENP-B boxes adjacent neocentromeric sites may be random. Alternative, their positioning could have functionality in making the chromatin state primed for neocentromere formation, while centromere formation is unlikely to enable “recruitment” of CENP-B boxes. This however cannot be completely excluded as CENP-B is an ancient *pogo*-like transposase and these may be recognition sequences that retain jumping skills, with inverted pairs of boxes best candidates for residual transposition activity.

This is in line with their enrichment at meiotic recombination sites, particularly intriguing since transposons preferentially insert at recombination hotspots and meiosis is when transposons are most active. Interestingly, overlaps between ECSs and DMC1 were detected across multiple chromosomes, but a preferential association was observed within the PARs, where meiotic exchange is obligatory. The absence of enrichment outside the PAR indicates that ECSs are not generally associated with recombination hotspots across the genome. This context-specific enrichment suggests that ECSs may acquire specialized functions in highly recombinogenic chromatin, potentially linking centromere-related sequence features with the regulatory landscape of meiotic recombination. While these potential roles remain speculative and will require further functional validation, our detailed classification represents a first step toward unraveling the complexity and biological significance of these newly identified genomic elements, whose functional relevance may extend well beyond their selected occurrence and basal structural roles especially in genomic contexts characterized by dynamic chromatin states.

## Conclusion

In summary, this work establishes that ECSs are not genomic relics but conserved elements with specific structural and organizational features whose activity depends on their chromatin environment. Our findings demonstrate that different subsets of ECSs, which we classified as “*ectocentromeric sites*”, can influence chromatin organization and gene expression, also through CENP-B recruitment in absence of underlying CENP-B boxes, and may also participate in broader processes such as meiotic recombination, chromatin looping and neocentromere containment. Together, these results redefine the ECSs as a new class of centromeric-like sites with a potential broad functional relevance, expanding the genomic reach of centromeric proteins and opening new directions for understanding chromosome organization and stability.

## Acknowledgments.

We thank Luca Corda for his key computational insights into the analysis, Alessio Colantoni for providing feedback during the project, and Michele Marass for editorial advice.

## Author contributions

S.G. conceived and supervised the study. P.S. carried out the analyses and generated figures. C.B. and F.B processed Hi-C sequencing data, including read alignment, contact matrix generation, and TAD calling. S.G. and P.S. wrote the paper.

## Supplementary Figure Legends

**Figure S1.**
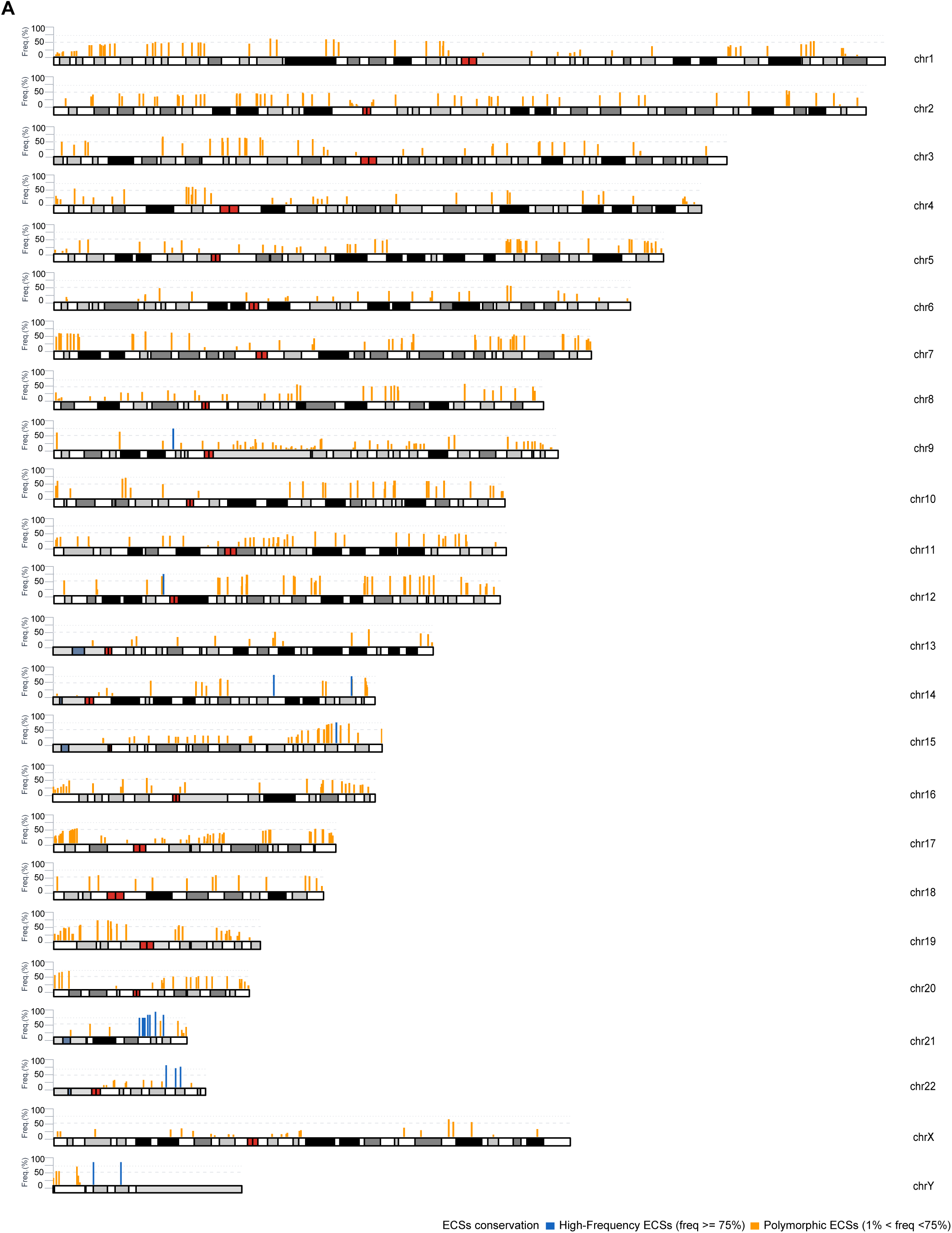
Karyotype representation of 854 CHM13v2.0 of ECSs with frequency >0% across 215 HPRC/HGSVC phased haplotypes from 109 individuals. Each bar represents one ECS position (width = 500 kb, centered on the site midpoint). Bar height is proportional to ECS frequency across the analyzed haplotypes (0–100%). Blue bars (n=18): High-Frequency ECSs with frequency ≥75%. Orange bars (n=836): Polymorphic ECSs with frequency between 0% and 74%. Chromosomal lengths reflect CHM13 T2T v2.0 coordinates. Dashed reference lines indicate 25%, 50%, and 75% frequency levels.

**Figure S2.**
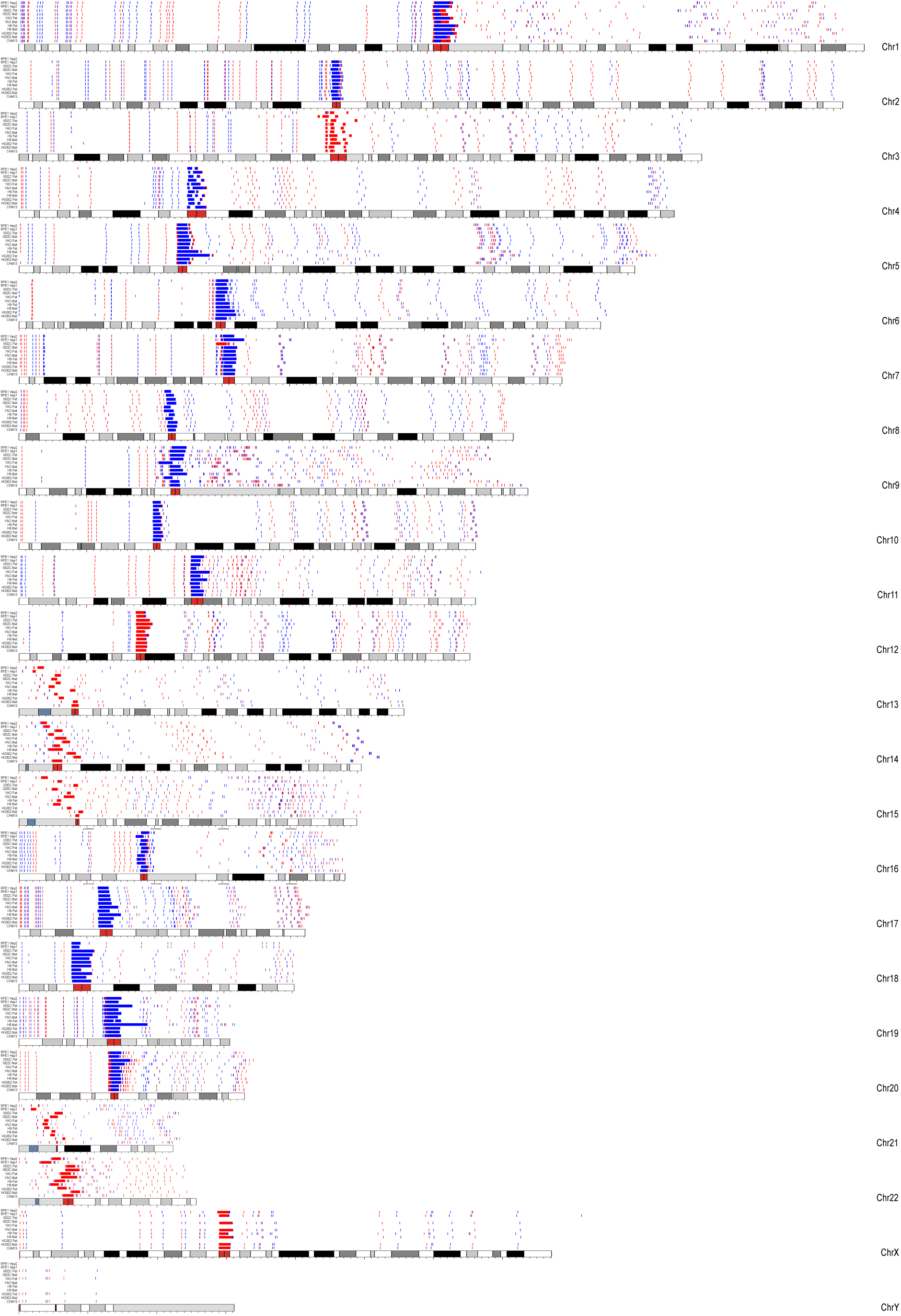
The Human Centeny Map across several T2T human genome assemblies. Blue depicts the CENP-B box in forward, while red represents its reverse complement within each genome. From the top to the bottom line, the genomes used for the analysis are: RPE-1v1.1 hap2 v1.0, RPE-1 hap1 v1.0, I002C Pat, I002C Mat, YAO Pat, YAO Mat, H9 Pat, H9 Mat, HG002 Pat, HG002 Mat, and CHM13v2. CHM13 ideograms from the *karyoploteR* (v1.28.0) package [55] were used for all chromosomes.

**Figure S3.**
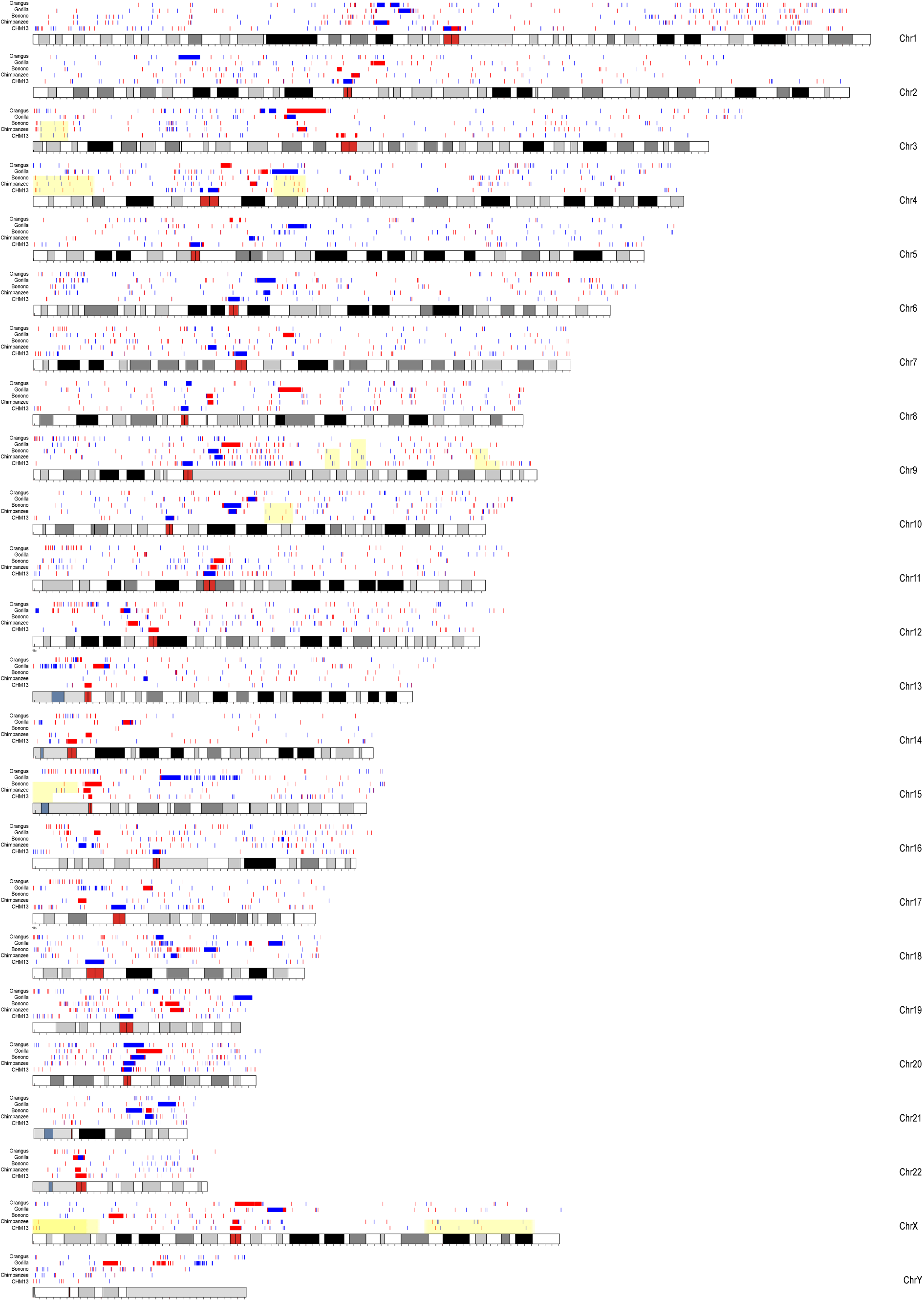
The “Inter-species Centeny Map” across several of the latest T2T primate genome assemblies. Blue depicts the CENP-B box in forward, while red represents its reverse complement within each genome. From the top to the bottom line, the genomes used for the analysis are: *Sumatran orangutan* (pongo abelii), *Gorilla gorilla gorilla* (western lowland gorilla), *Pan paniscus* (pygmy chimpanzee), and *Pan troglodytes* (chimpanzee). The regions highlighted in yellow indicate a similar ECS pattern between primate and human genomes. CHM13 ideograms from the *karyoploteR* (v1.28.0) package [55] were used for all chromosomes.

**Figure S4.**
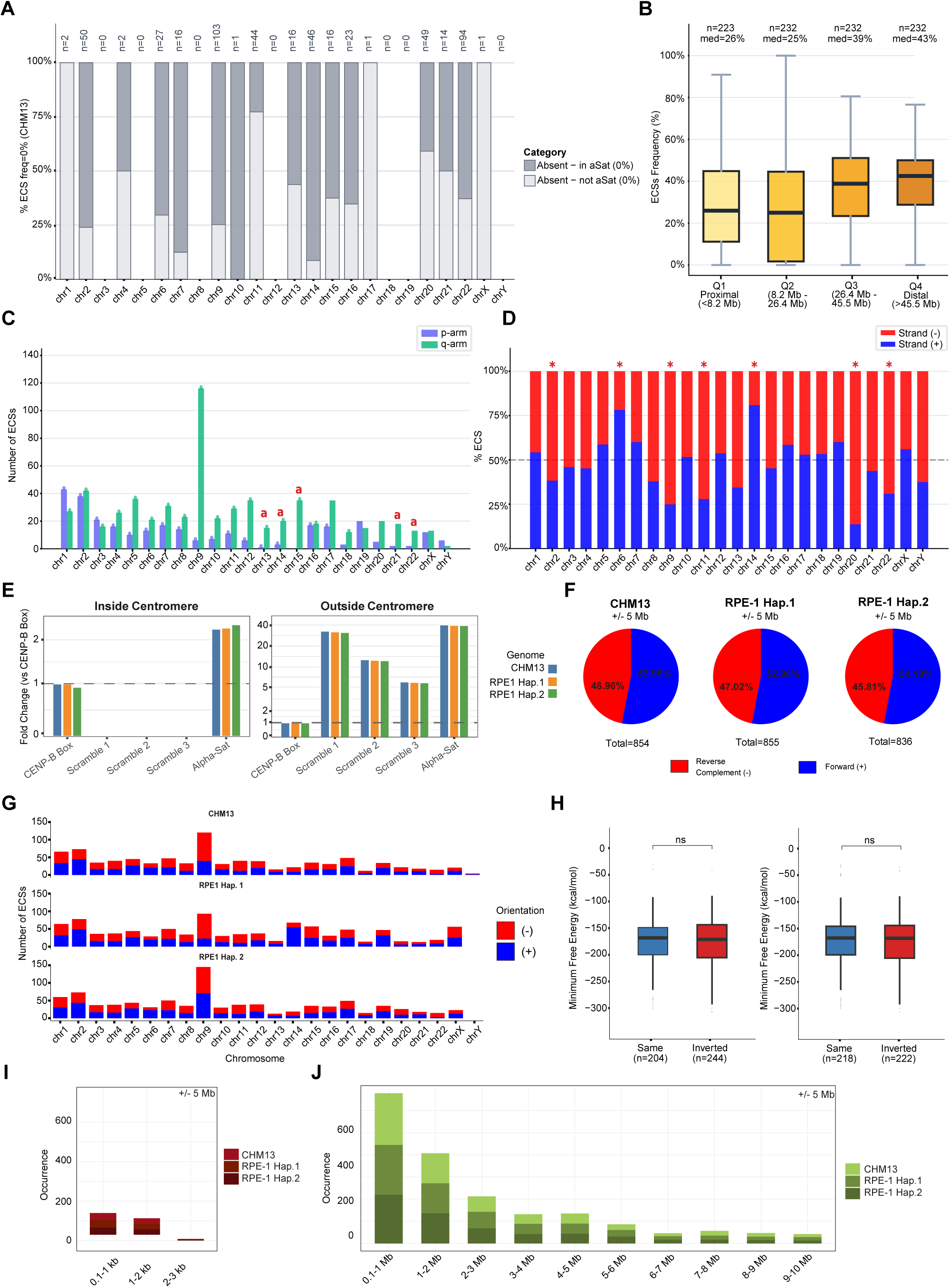
A) Stacked bar chart showing the distribution of 505 CHM13-specific ECS positions (freq = 0%) across HPRC/HGSVC haplotypes. Most absent ECS overlap with α-satellite regions (63.8%), likely reflecting structural variability of pericentromeric HOR arrays (dark grey), while the remaining sites (36.2%) fall outside α-satellite regions and likely represent non-functional motif matches (light grey). Numbers above bars indicate the total number of absent ECS per chromosome. B) Box plots of ECS frequency (%) grouped by quartile of distance from the centromeric border. Q1 = proximal (<8.2 Mb from centromere); Q2 = 8.2–26.4 Mb; Q3 = 26.4– 45.5 Mb; Q4 = distal (>45.5 Mb). C) Bar chart showing the number of ECS positions per chromosomal arm (p arm, purple; q arm, green) for each chromosome. D) Stacked bar chart showing strand orientation bias (strand+, blue; strand−, red) per chromosome. Asterisks () indicate significant deviation from the expected 50/50 ratio (chi-square test, p<0.05). E) Bar plot showing the fold change of CENP-B Box, scrambled, and αConsensus sequences, either inside or outside the centromere, in both CHM13 and RPE-1 genomes. F) Pie charts representing the genome-wide total counts and the percentages of forward and reverse complement ECSs in CHM13 and RPE-1 haplotypes 1 and 2 after applying a 5 Mb cut-off. For the analysis, we considered the occurrences within the divergent αSat arrays located outside the centromere, provided by HumAS-HMMER_for_AnVIL. G) Stacked bar plots showing the total number of forward and reverse-complement ECSs per individual chromosome following the application of a 5 Mb cut-off. H) Boxplot of Minimum Free Energy (MFE) distributions for sequences flanking ECS motifs considering motif orientation. I-J) Stacked bar plots represent the detailed analysis of the (I) short- and (J) broad-range groups of ECSs distance values. H) Bar plots showing the distribution of ECS colocalizing with the ATAC-Seq peaks.

**Figure S5.**
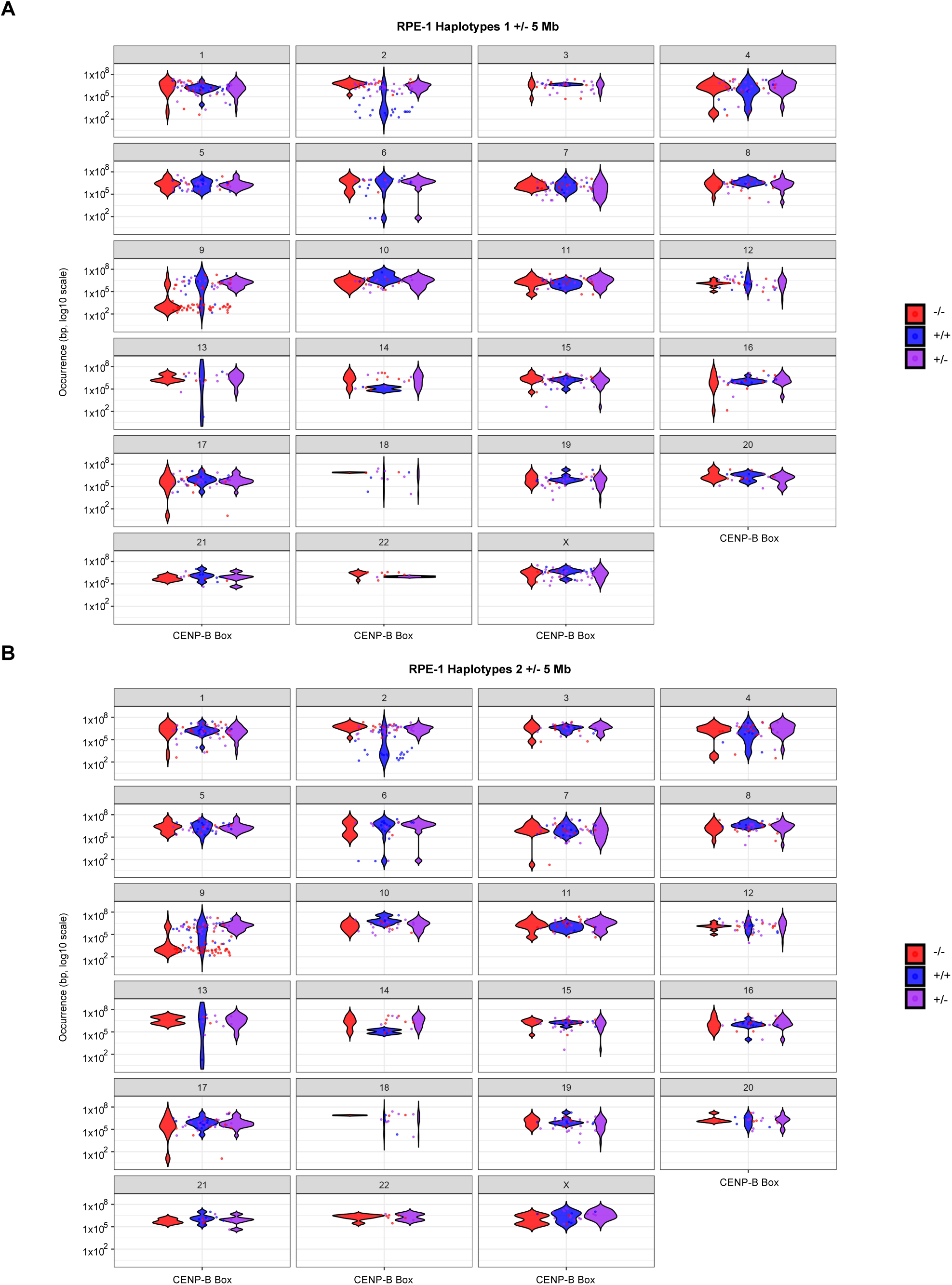
Violin plots depicting the distributions of ECS distance values across all chromosomes in RPE-1v1.1 haplotype 1 (A) and haplotype 2 (B), analyzed after applying a 5 Mb cut-off.

**Figure S6.**
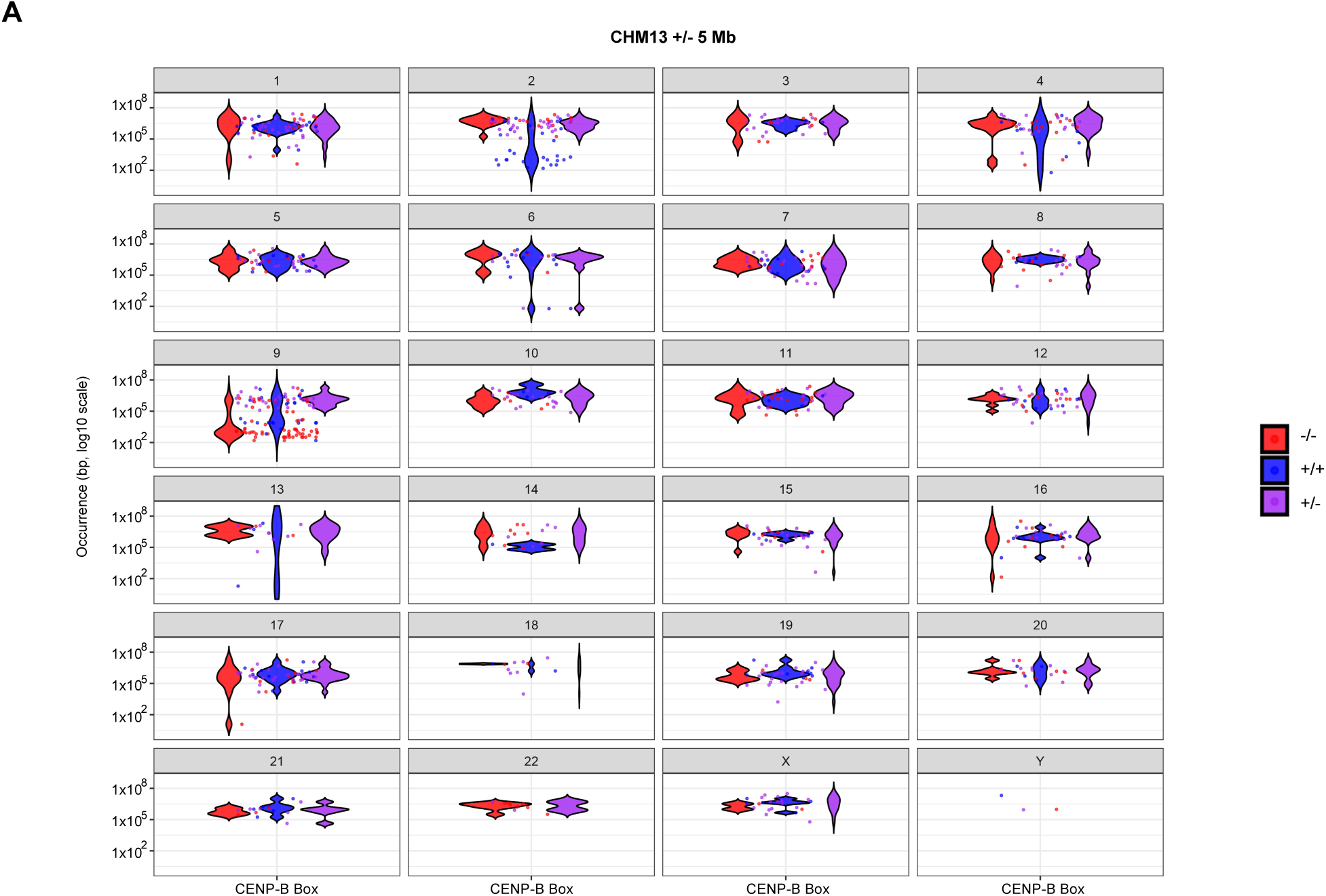
Violin plots depicting the distributions of ECS distance values across all chromosomes in CHM13v2 (A), analyzed after applying a 5 Mb cut-off.

**Figure S7.**
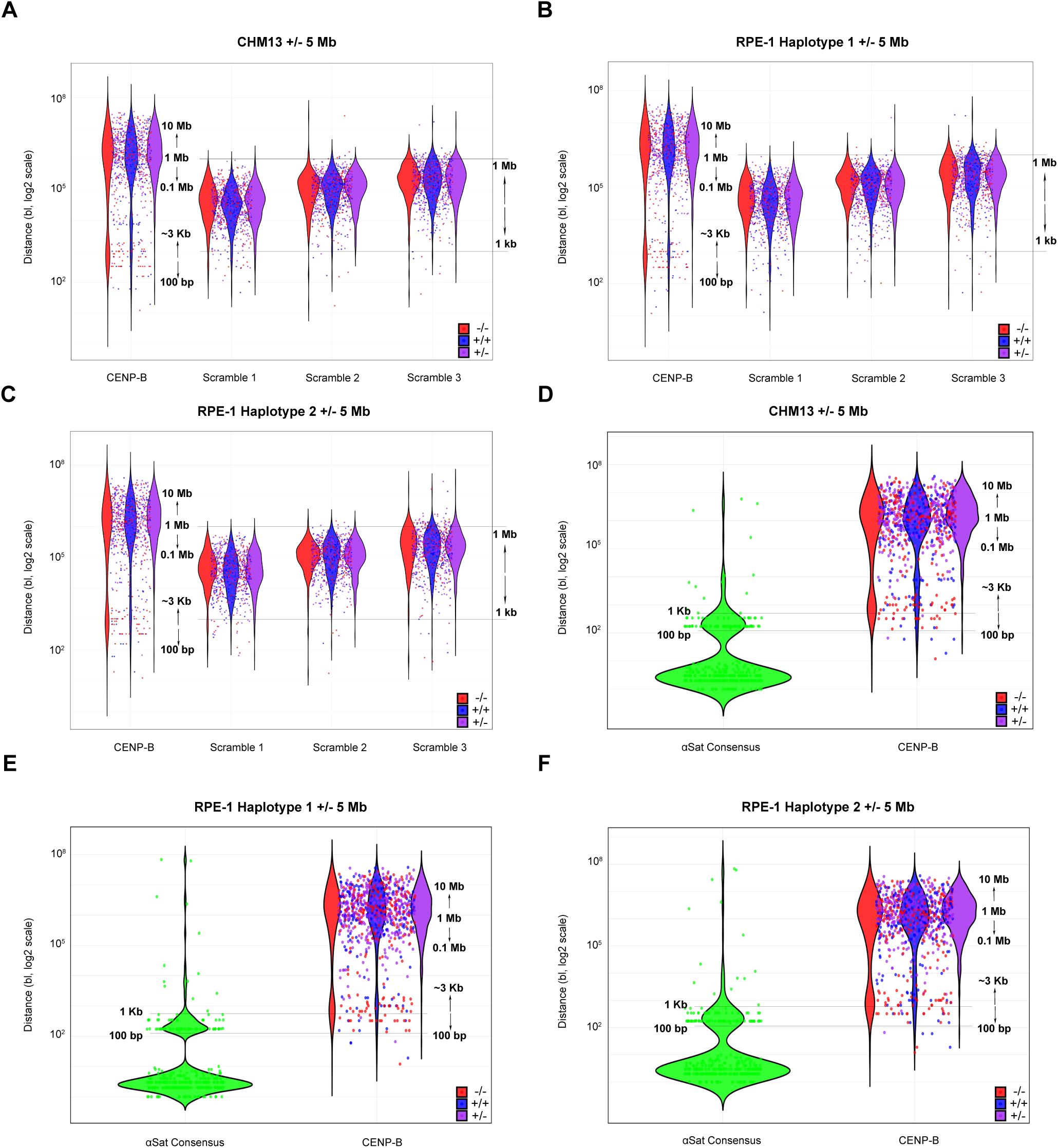
Violin plots represent the genome-wide distance value distributions of ECSs compared with those of the three scrambled (A-C) and the αConsensus (D-F) sequences generated in both CHM13v2 and RPE-1v1.1 reference genomes, analyzed after applying a 5 Mb cut-off.

**Figure S8.**
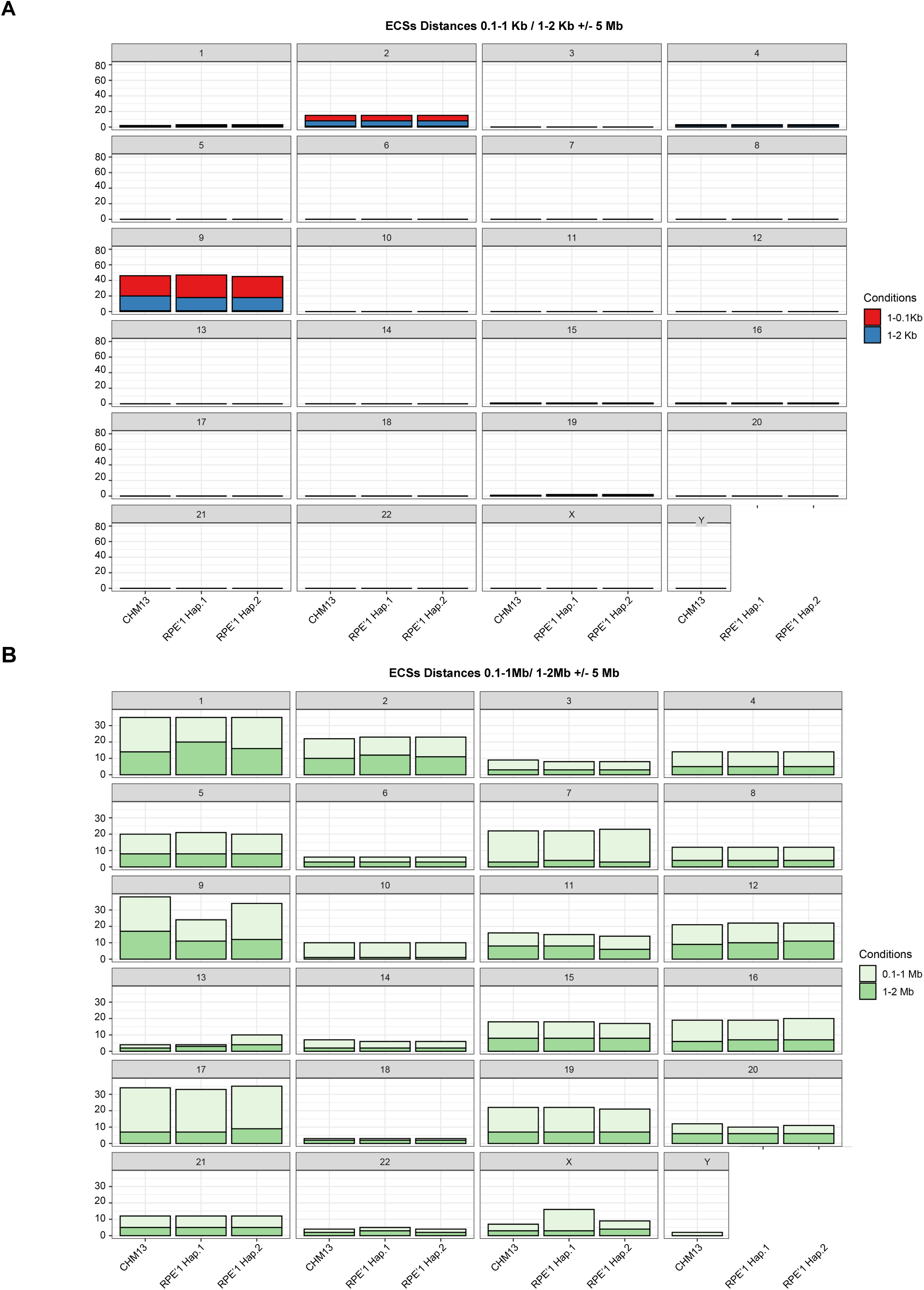
Bar plots showing the most representative ECS distance values across all chromosomes in the long-range group (0.1–1 Mb and 1–2 Mb; A) and the short-range group (0.1–1 kb and 1–2 kb; B) in the CHM13 and RPE-1 haplotype 1 and 2 genomes, after applying the 5 Mb centromeric exclusion.

**Figure S9.**
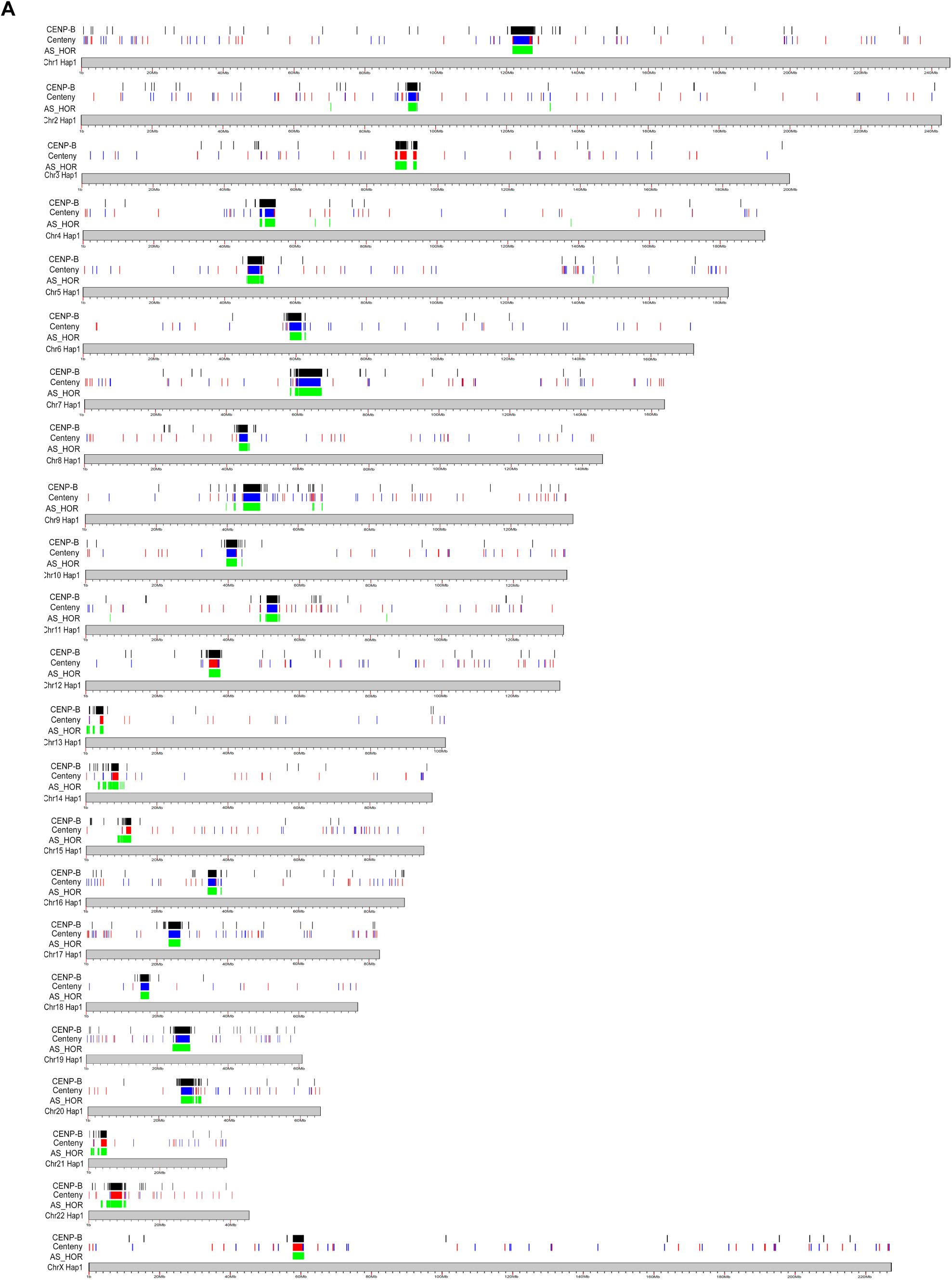
The human centeny map of the ECS motifs along p- and q-arms and centromeres of all chromosomes for the RPE-1v1.1 haplotype 1 genome assembly compared to CENP-B protein peaks and αSat (AS-HOR) annotation.

**Figure S10.**
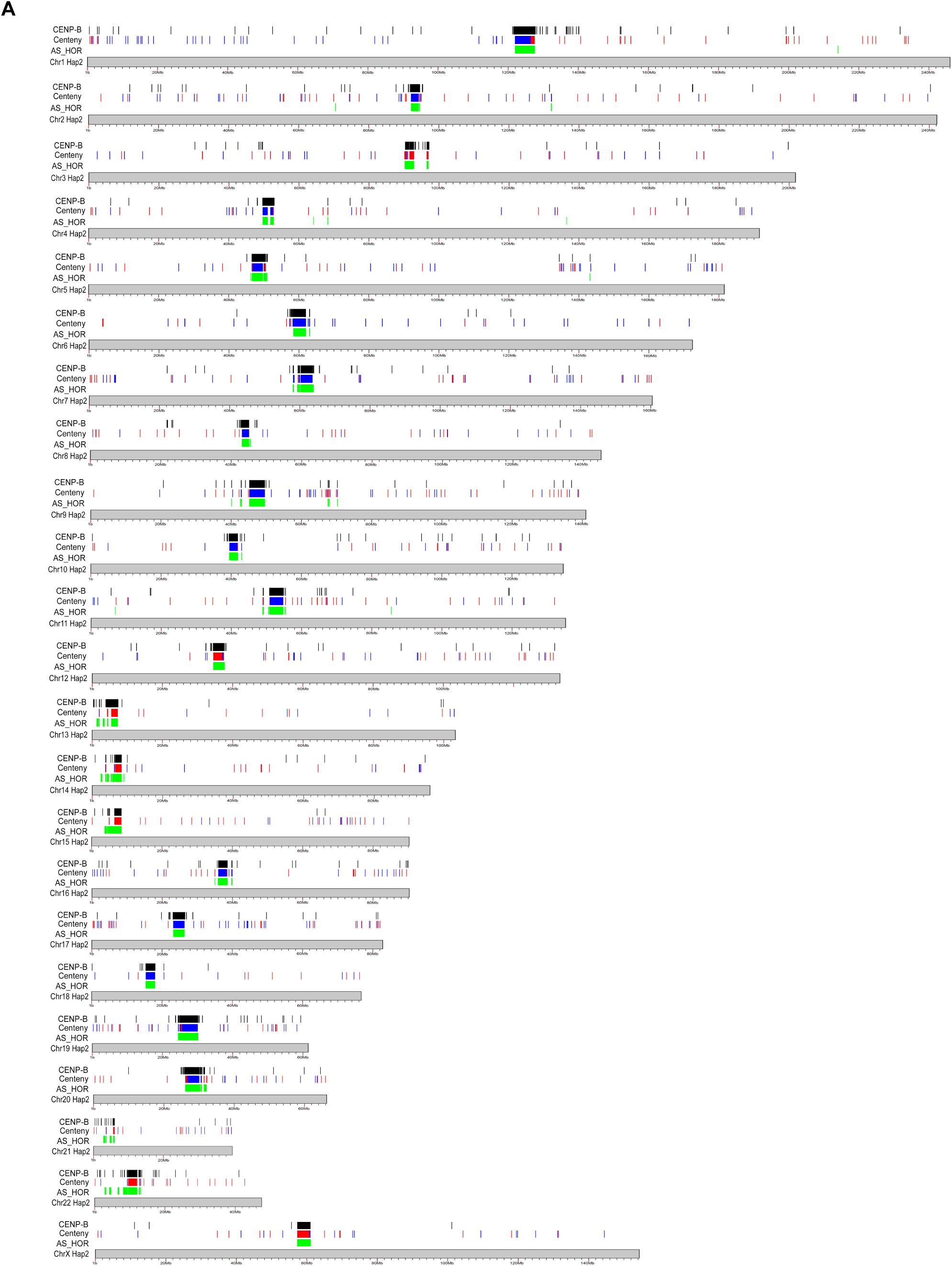
The human centeny map of the ECS motifs along p- and q-arms and centromeres of all chromosomes for the RPE-1v1.1 haplotype 2 genome assembly compared to CENP-B protein peaks and αSat (AS-HOR) annotation.

**Figure S11.**
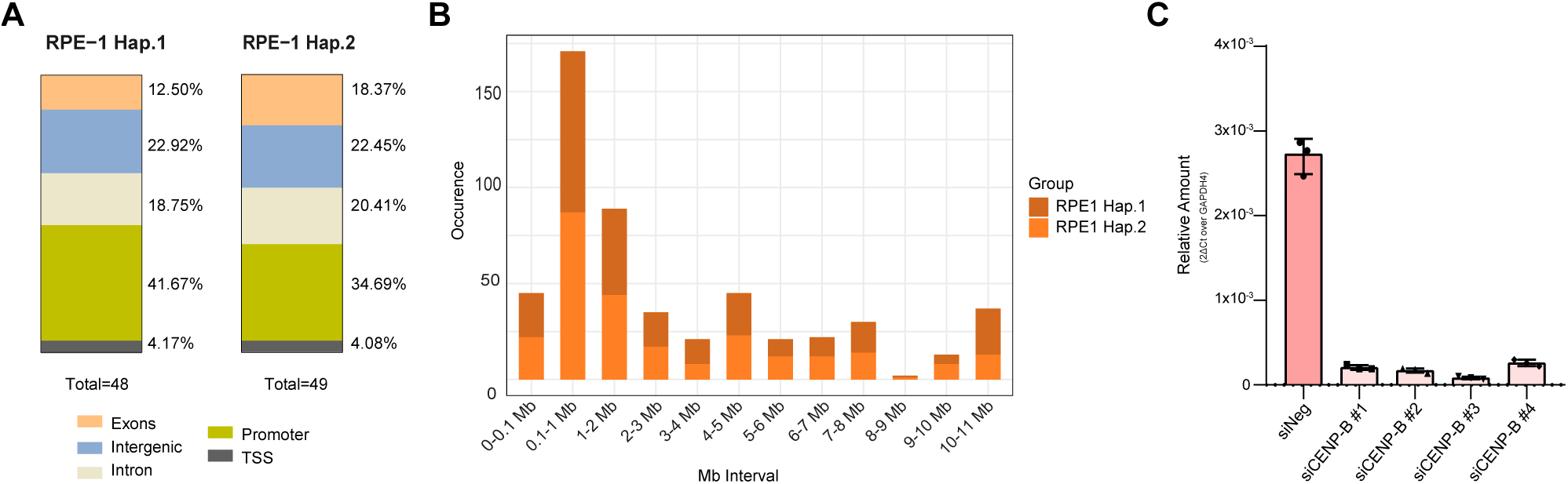
A) Bar plots showing the distribution of ECS colocalizing with the ATAC-Seq peaks. B) Stacked bar plot showing the number of CENP-B binding sites in the box–near-protein category colocalizing with ATAC-seq peaks at defined distance ranges from the nearest ECS. C) RT–qPCR analysis of four different siRNAs targeting CENP-B.

**Figure S12.**
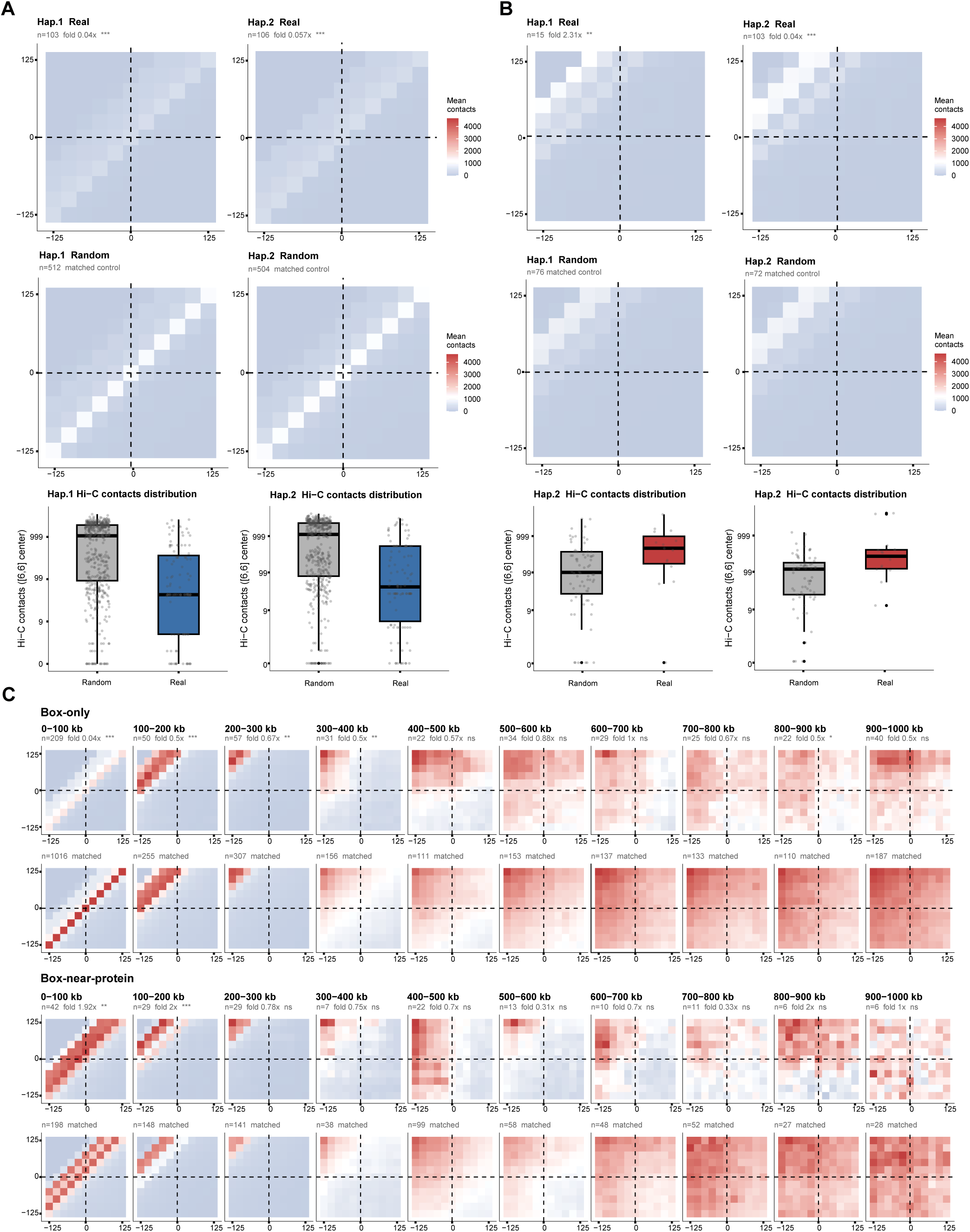
A) APA heatmaps and boxplots showing Box-only pairs at 0–100 kb, analyzed separately per haplotype (top, haplotype 1; bottom, haplotype 2). For each haplotype: mean real APA, mean distance-matched random APA, and boxplot of central [6,6] contact values (log10) for real versus random pairs. B) APA heatmaps and boxplots showing Box-near-protein pairs at 100–200 kb, analyzed separately per haplotype (top, haplotype 1; bottom, haplotype 2). For each haplotype: mean real APA, mean distance-matched random APA, and boxplot of central [6,6] contact values (log10) for real versus random pairs. C) APA heatmaps (11×11 bin, ±125 kb, 25 kb resolution) for Box-only and Box-near-protein pairs across all 10 distance bins (from 0–100 to 900–1000 kb), arranged in 4 rows: Box-only real, Box-only random, Box-near-protein real, Box-near-protein random (Hap1 + Hap2 pooled). Color scale calibrated independently per bin. Fold-change and Wilcoxon significance are annotated below each real panel. Box-only depletion extends across the first 4 bins (0–400 kb); Box-near-protein enrichment is confined to the first 2 bins (0–200 kb), demonstrating the distance specificity of the double dissociation.

